# Historical biogeography and population genetic structure of the giant gilded catfish (*Brachyplatystoma rousseauxii*): expanding Humboldtian connectivity routes between the Orinoco and Amazon River basins

**DOI:** 10.64898/2026.08.04.742678

**Authors:** José Gregorio Martínez, Diana Sanchez-Bernal, Sandra Hernández-Rangel, Jacqueline Batista, Susana J. Caballero, Izeni Pires Farias, Tomas Hrbek

## Abstract

Understanding the evolutionary history of species within a geographic context is key to historical biogeography, as it reveals how geological and climatic changes shaped biodiversity. This is especially important in ecologically significant regions like the Amazon and Orinoco basins. Together, they host the world’s greatest freshwater fish diversity (∼3,500 species), sharing a common but not yet fully understood evolutionary history. The gilded catfish (*Brachyplatystoma rousseauxii*), an ancient species widely distributed as a metapopulation in Neotropics, is an important model for studying past connectivity, divergence, and historical processes shaping fish diversity between these basins. This study analyzed the genetic structure, connectivity routes, and demographic history of *B. rousseauxii* using nuclear (microsatellite and ddRADseq) and mitochondrial DNA. Population structure analyses and coalescent models indicate that *B. rousseauxii* populations from the Orinoco and Amazon basins are genetically distinct, with no evidence of current gene flow. However, our results support the occurrence of a possible secondary contact event after the divergence, with the Boa Vista population retaining the genetic signal of this process. The ancestral population split occurred at the Rupununi Portal around 2.54 Ma (ddRAD) or 1.31 Ma (mtDNA). Then, the species colonized the Branco and Orinoco Rivers ∼1.90 Ma (ddRAD) or 0.6 Ma (mtDNA), rapidly expanding in the Orinoco (>1.3 or >0.29 Ma), while colonization of the Amazon from the Branco River was more recent (≤1.0 or ≤0.15 Ma). Population expansion signal was detected in the Orinoco (∼0.20 Ma), whereas the Amazon remained stable. Our findings suggest that the rise of the Vaupés Arch in the Late Miocene does not explain the observed genetic divergence. Likewise, the Casiquiare Canal and Japurá-Guaviare headwaters are not connectivity routes between basins. Instead, the Rupununi Portal, including the recent capture of the Branco River by the Negro River, was the last point of connection and played a key role in shaping *B. rousseauxii*’s distribution. These findings provide insights into Neotropical fish biogeography and the historical configuration of the Orinoco and Amazon basins.

## Introduction

One of the main objectives of modern historical biogeography is the reconstruction of the evolutionary history of a particular species or groups of species in a geographic context, to understand how geological and climatic changes influenced dispersal, migratory events, divergence, range expansion, and demographic events in a particular area or ecoregion [1–5].

In Neotropics, the Amazon and Orinoco are respectively the second and third longest rivers in the world with approximately 6437 and 2140 km in length, respectively [6,7]. Both basins are characterized by their high ichthyofaunal diversity, estimated to be around 3500 fish species [4], with almost 65% of all neotropical ichthyofauna concentrated in their waters [8]. Much of this diversity is shared between these two basins [4], nonetheless, little is known regarding the extent of current and historical biological, demographic, and reproductive connectivity between the fish faunas of these basins. However, three contemporary routes of ichthyofaunal connectivity between these basins have been hypothesized: (i) the Casiquiare Canal, currently the most evident and best-supported connection; (ii) the headwater connection between the white-water Guaviare River (Orinoco basin) and the Japurá River (Amazon basin) on the Andean piedmont; and (iii) the Rupununi Portal in the Guiana Shield region (see connectivity hypotheses in Fig 1).

**Fig 1.**
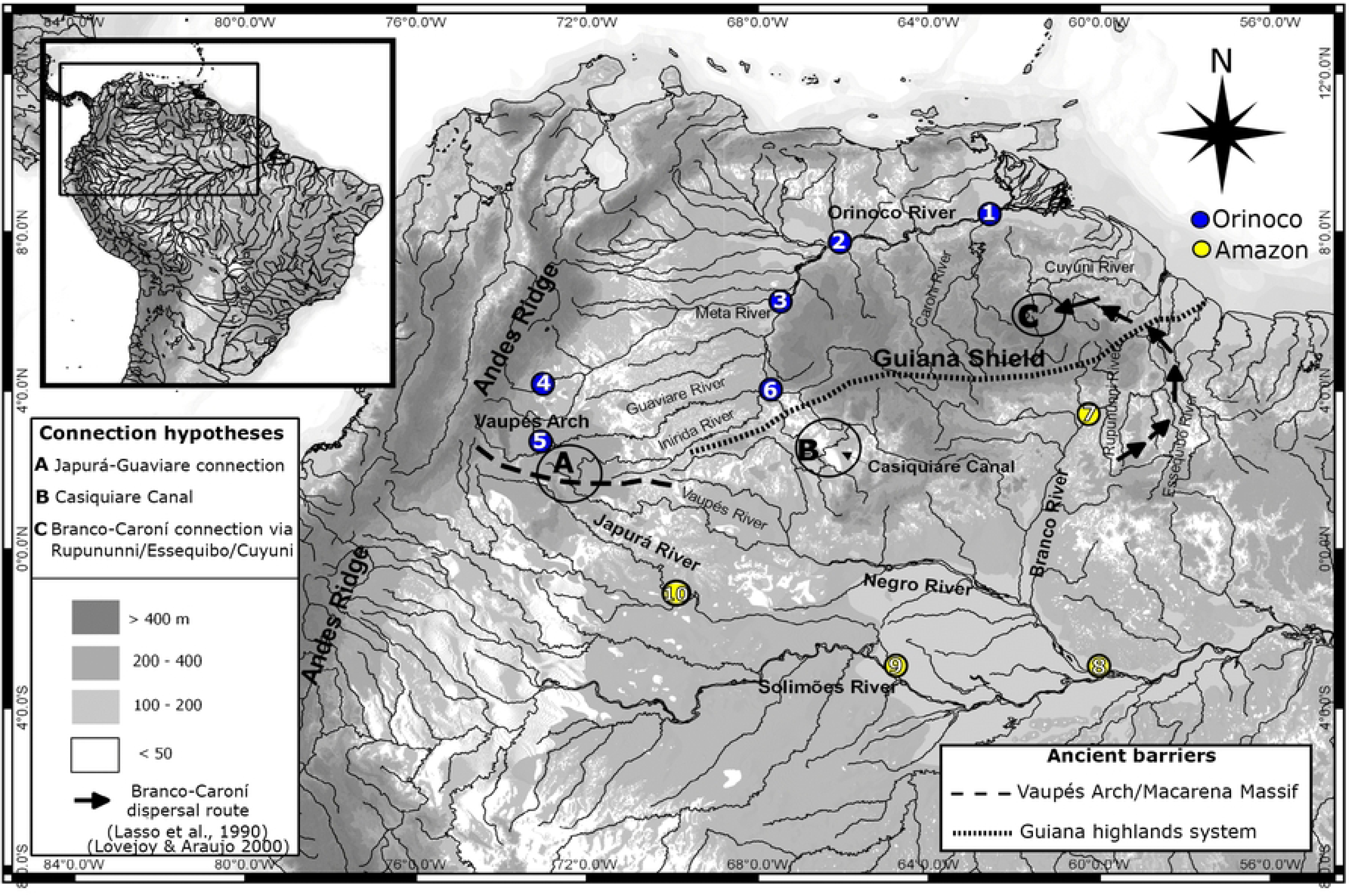
Map of the hydro-geological scenario and study of sampling sites. Sampling localities for the gilded catfish *Brachyplatystoma rousseauxii* individuals from the Orinoco [Ciudad Guayana (1), Caicara del Orinoco (2), Puerto Carreño (3), Puerto López (4), Guaviare (5), Inírida (6)] and Amazon basin [Boa Vista (7), Manaos (8), Tefé (9), La Pedrera (10)]. Points A, B and C represent the three hypotheticals connectivity routes to test for the gilded catfish between the Orinoco and Amazon basins. River network and background raster data were obtained exclusively from Natural Earth (www.naturalearthdata.com), which provides publicly available geographic datasets licensed under the Creative Commons Attribution 4.0 (CC BY 4.0). No copyrighted or proprietary data sources (e.g., Google Maps, Google Earth, or similar platforms) were used in the creation of this figure. The map was constructed using these datasets within the R statistical environment (www.r-project.org) and edited using Inkscape v1.X (https://inkscape.org).

Physical connectivity between these two basins was unknown to science until the discovery of the Casiquiare Canal in 1799, when the naturalist Alexander Von Humboldt explored the Orinoco and Amazon Rivers, providing the first scientific evidence of the existence of this connection. This discovery was somehow expected, considering the common historical geological origin of both basins. Hoorn et al. (1995) [9] and Lundberg et al. (1998) [10] presented evidence showing that the western Amazon River had been connected to the Orinoco, forming a unique transcontinental basin called the Paleo-Orinoco-Amazonas from 25 Ma until approximately 11-13 Ma, when the Vaupés Arch emerged, extending itself from the Andes to the Guiana Shield, separating the two basins. Regardless of the separation of both basins in the Miocene, a fluvial connection persists between them. The most predominant is the Casiquiare Canal, an arm of the upper Orinoco River that flows into the Negro River, the main affluent of the Amazon River. The Casiquiare is a black water river, with acidic waters and a length of 300 km, approximately width of 100 m at the source, and an average discharge of 2100 m^3^/s, flowing closer to the southwestern border of the Guiana shield and the western portion of the “Llanos” (savannas) of the Orinoco River in its contact zone of the Western Amazonia [11]. The Casiquiare substantially increases its length and volume during its course, since it receives water from tributaries such as the rivers Pamoni, Pasiba, Siapa, and Pasimoni. In its mouth, of approximately 500 m in width, it joins the Guania River to form the Negro River, the main tributary of the Amazon River [11].

The physical connectivity of the Casiquiare is, up to date, the most frequent hypothesis used to explain the observed patterns of current and historical fish dispersion between the Amazon and Orinoco basins [5,10–14]. However, the Casiquiare could also act as an ecological filter, mainly due to its strong pH gradient along its course. For many species, the Casiquiare becomes a barrier to dispersion [11] or, at least, it strongly limits gene flow (*e.g*. species *Cichla* [5]; ocellate freshwater stingray, *Potamotrygon motoro* [15]).

The Casiquiare Canal is not the only aquatic connection between the upper Orinoco and the Negro River; at least five alternative connections between affluents of the Orinoco and Negro towards the east of the Casiquiare Canal [14,16]: Guacamayo creek, right bank affluents of the Conochirite River and tributaries of the Atacavi and Temi Rivers (Orinoco basin), floodplains of the Baria River (affluent of the Pasimoni River, Casiquiare canal drainage) and the Maturaca River (affluent of the Cauaburi River, Negro River drainage), and finally, via the Pimichin creek, a connection already mentioned by Alexander von Humboldt and confirmed by Rice (1921) [17].

The Amazon and Orinoco Basins are also connected by the headwaters of the white-water Rivers Guaviare (Orinoco) and Japurá (Amazonas) on the Andean piedmont during flooding periods, suggesting an alternative hypothesis of a main corridor in the Western Amazon, particularly for migrant species, such as big catfish species (Lundberg pers. com.).

Finally, a connection between the two basins via the Rupununi portal has also been suggested. This connection was initially suggested for the sedentary fishes of the genus *Potamorrhaphis* [18], based on analysis of mtDNA phylogeographic patterns. A recent phylogeographic study of *Piaractus brachypomus*, a species distributed across both basins and characterized by regional reproductive migrations, suggested that historical connectivity via the Rupununi Portal was more likely than through the Casiquiare Canal, based on the patterns of shared ancestry observed between basins near this region [19].

The Rupununi portal, on the Guiana Shield, is localized in southern Guyana, on uplands at > 300 m.a.s.l. [20]. This portal is a unique biogeographic area allowing a seasonal connection between the Amazon River (via the Takutu/Branco/Negro Rivers) and the Essequibo River (via the Rupununi River). Guiana rivers are recognized as biogeographic areas of endemism for many taxa [21–23]. This hydrological connection occurs during the annual rainy season from the inundation of the Rupununi savannas, encompassing about 3,480 km^2^ with a hydric period of 49 days [24], allowing for ichthyofaunal exchange between them [18,25–27]. However, it could also function as a barrier for dispersal for some species.

It is unclear which of these areas, Japura/Guaviare headwaters (Andean piedmonts), Orinoco/Negro headwaters (Casiquiare canal and floodplains) or the Rupununi portal (Orinoco mouth and Branco River), most likely served and/or serves as a conduit between the ichthyofaunas of the two basins, or if the division of the Paleo-Orinoco-Amazonas basin into the Amazon and Orinoco basins resulted in vicariant divergence of the faunas of the two basins.

The difficulty of testing these connectivity alternatives has been due mainly to the lack of suitable data to test these alternate hypotheses. To test these hypotheses we collected mitochondrial, microsatellite and genomic data from the gilded catfish (*Brachyplatystoma rousseauxii*). *Brachyplatystoma rousseauxii* is an apex predator and one of the largest South American fish (up to 192 cm) migrating up to 5500 km for reproduction during 5–6 months [12,28]. It is also an important commercial species [12,28–31] and is widely distributed in the Orinoco and Amazon basins. Beyond its remarkable migratory capacity, the ecology and life history of *B. rousseauxii* make it a biologically plausible candidate for inter-basin dispersal. The species is strictly associated with nutrient-rich white- and clearwater drainages [12,28], making the blackwater Casiquiare Canal (pH 2.9–4.2; conductivity ∼8 µS) a challenging hydrochemical barrier, while clearwater corridors such as the Branco River tributaries and the Rupununi Portal probably offer more hospitable routes. Adults undertake the longest migration recorded for any strictly freshwater fish — up to 12,000 km round trip to spawning grounds in the Andean foothills (170–280 m asl) [12,27,32] — enabling them to reach remote headwater areas near inter-basin water divides inaccessible to less vagile species. Larvae then drift passively 4,000–5,500 km downstream to estuarine nurseries [12,27,32], meaning reproductive events near headwater connections could passively transport ichthyoplankton into adjacent drainages. Natal homing demonstrated by otolith strontium analysis [33,34] imposes population fidelity to natal rivers, yet also implies that rare straying events at headwater junctions can establish founder populations in neighboring basins. The life cycle of *B. rousseauxii* is thus entirely dependent on longitudinal river connectivity [12,27,32], making it exquisitely sensitive to the barriers and corridors created by geological and hydrological reorganization.

The aim of this study was to reconstruct the historical biogeography and evaluate contemporary demographic connectivity of *Brachyplatystoma rousseauxii* between the Orinoco and Amazon basins using mitochondrial, microsatellite, and genomic data. Specifically, we investigated patterns of genetic structure, historical and contemporary gene flow, phylogeographic relationships, and demographic history across both basins. We further tested whether the divergence observed between populations is better explained by historical vicariance associated with the fragmentation of the Paleo-Orinoco- Amazonas system following the uplift of the Vaupés Arch, or alternatively by historical and/or contemporary connectivity among basins through currently hypothesized dispersal routes.

## Materials and methods

### Sample and data collection

Tissue samples were collected from six localities in the Orinoco basin [Ciudad Guayana (1), Caicara del Orinoco (2), Puerto Carreño (3), Puerto López (4), Guaviare (5), Inírida (6)] and four localities in the Amazon basin [Boa Vista (7), Manaos (8), Tefé (9), La Pedrera (10)] (Fig 1), totaling 154 samples, 73 from Orinoco and 81 from Amazon (S1 Table). In Brazil, permission to collect tissue samples was granted by SISBIO/IBAMA (permit: #49641-2), and in Colombia by ANLA (Permit: Resolution 1177 (09 Oct 2014)) to Universidad de Los Andes.

This study does not involve human participants, human biological samples, or identifiable personal data. The research focuses exclusively on a native freshwater fish species and its population genetic and biogeographic patterns. Tissue samples were obtained from specimens acquired at municipal fish markets in each sampled city; these specimens originated from artisanal fisheries operating legally within the respective river basins. No animals were captured or sacrificed specifically for this study.

Therefore, approval by an animal research ethics committee was not required. All tissue samples were deposited in the Laboratory of Evolution and Animal Genetics (LEGAL) at the Universidade Federal do Amazonas (S1 Table).

Tissue samples were collected from whole, intact individuals at artisanal fish landing sites, arriving at sampling points at approximately 06:00 h, prior to any fish processing or filleting, so that all diagnostic morphological characters remained available. Species identity was confirmed through a multi-step verification protocol. First, *B. rousseauxii* was identified by its unique silvery-platinum head and characteristic golden-yellow body coloration—a combination absent in all congeners, including *B. vaillantii*, *B. filamentosum*, and *B. platynemum*—following the diagnostic criteria of Lundberg & Littmann (2003) [35]. This morphological assessment was cross-validated with local artisanal fishers, whose traditional ecological knowledge of the species (*dourada/dorado*) constitutes a reliable complementary criterion given the commercial importance of this taxon across the fisheries of all Orinoco and Amazonian countries where it occurs. Definitive species confirmation was provided by the COI barcode sequences generated for this study: of the 154 samples collected under those criteria, 82 were randomly selected for phylogeographic analyses and molecular identification, ensuring representative coverage of all sampling localities within each basin. Species identity was subsequently verified through BLASTn searches against GenBank databases, and only sequences exhibiting ≥98% identity to vouchered *B. rousseauxii* reference sequences were confirmed and retained (S1 Table, Sheet 2), a threshold widely accepted for unambiguous fish species identification [36,37]. Additionally, pairwise Kimura 2-parameter (K2P) genetic distances among all COI sequences estimated in MEGA X [32] within (Orinoco: 0.09%; Amazon: 0.01%) and between basins (0.24%) did not exceed 2%—the established intraspecific threshold in fish barcoding [36]—while interspecific divergence within *Brachyplatystoma* ranges from 2.1% to 8.3% at COI [38], making the inadvertent inclusion of misidentified congeneric individuals readily detectable and effectively excluded by this criterion.

DNA extraction was performed using standard phenol chloroform protocol [39,40]. Quality of the extraction was evaluated using 0.8% agarose gel stained with GelRed (Biotium). Quantification of DNA was determined spectrophotometrically by Nanodrop 2000 Thermo-Scientific and diluted to a final concentration of 50 ng/uL.

To obtain genetic data, both nuclear and mitochondrial markers were amplified. Data included microsatellite genotypes (n=154), using the primers developed by Batista et al. (2009) [41]; analysis of SNPs obtained by *de novo* genotyping on IonTorrent PGM (n=30), following Martínez et al. (2016) [42]; sequencing of the COI marker (n=82), using the universal cocktail primers developed by Ivanova et al. (2007) [43]; and finally, sequencing of the mitochondrial (mtDNA) control region (n=82), according to Sivasundar et al. (2001) [44]. For each marker, homogeneous sample representation was used for each basin. For the population structure analyses, due to the low sample size (n<4 individuals), samples from Inírida location were grouped with samples from the Guaviare location; and the samples from Caicara del Orinoco, were grouped with Guayana.

PCR amplification conditions for microsatellites (BR 7, BR 37, BR 38, BR 40, BR 41, BR 44, BR 45, BR 46, BR 47, BR 48, BR 49, BR 51) included a final volume of 10 µL containing 1µL of genomic DNA (50 ng/µL), each forward and M13 Label primer (FAM; 0.4 µM), reverse primer (0.8 µM), dNTP (10 mM), MgCl_2_ (1.5 mM), buffer (1X; 10 mM Tris–HCl, 50 mM KCl, pH 8.4) (Fermentas) and 0.5 U of Taq DNA Polymerase (1 U/μL, Fermentas). PCR temperature cycles program was conducted according to Batista et al. (2009) [41]. PCR products were genotyped in an automatic ABI 3500 sequencer. Fragment analysis was performed in the software GeneMapper. Allele sizes were inferred using the pUC19 ROX-labelled size standard [45]. Finally, as a result, a genotype matrix for each individual was constructed.

The SNPs development included construction of sequence libraries which were prepared using a variant of the ddRAD protocol [46] adapted for IonTorrent PGM platform (Life Technologies^TM^) and described in Martínez et al. (2016) [47], with Csp6I and SdaI restriction enzymes (Thermo Scientific). For more details, see the complete protocol on GitHub (https://github.com/legalLab).

The PCR mix for amplification of the mitochondrial DNA D-loop region was done in a final volume of 15 μL, with 5.1 μLof water, 1.5 μL of BSA (10 mg/mL), 1.2 μL MgCl_2_ (25 mM, Fermentas), 1.5 μL (NH_4_)_2_SO_4_ 10X buffer (Fermentas), 1.2 μL dNTPs (10 mM), 1.5 μL of each primer (2 μM), 0.5 μL *Taq* polymerase (1 U/μL, Fermentas) and 1.0 μL of DNA (∼ 50 ng/μL). For COI, the PCR mix was on a final volume of 15 μL containing 8.3 μL of water, 1.2 μL MgCl_2_ (25 mM, Fermentas), 1.5 μL (NH_4_)_2_SO_4_ 10X buffer (Fermentas), 1.2 μL dNTP (10 mM), 1.5 μL primer cocktail (a mixture of the *forward + reverse primers*) (2 μM), 0.3 μL *Taq* polymerase (1 U/μL, Fermentas) and 1.0 μL of DNA (∼ 50 ng/μL). All reactions were conducted for either COI or D-loop in a Veriti® (*Applied Biosystems*) thermocycler, following the amplification program: 72 °C for 1 min, 35 cycles of 94 °C for 10 sec, annealing temperature of 50 °C for 35 sec, 72 °C for 90 sec and a final extension step (72 for 5 min).

All successfully amplified products were checked on a 2% agarose gel and bidirectionally sequenced on a sequencer ABI-3500 (*Applied Biosystems*; www.appliedbiosystems.com) using the *BigDye Terminator Cycle Sequencing* v.3.0 kit (*Applied Biosystems*) chemistry.

### Data analysis

Mitochondrial sequences were checked and edited manually with Geneious v.10.0.9 [48]. The alignment to create consensus sequences was performed by using the integrated algorithm ClustalW [49]. Next, sequences were submitted to a BLAST search [50] for verification and comparison with sequences available in GenBank, and then checked manually for insertions, deletions and stop codons (COI) by using *in silico* translation in Geneious v.10.0.9. For phylogeographic inferences, COI and D-loop sequences were concatenated following Willis et al. (2010) [5]. These sequence data have been submitted to the GenBank database under accession number PX766138-PX766219 (D-loop) and PX769225-PX769306 (COI).

Genomic data quality was analyzed in FASTQC (Bioinformatics Group at the Babraham Institute, UK). The samples were demultiplexed using cutadapt [51], and SNPs were extracted using DiscoSnpRad [52] with default parameters except that presence indels and up to five SNPs per locus were permitted for construction of the loci, following the genotyping protocol of Escobar et al. (2024) [53]. The resulting Variant Call Format (VCF) was quality filtered considering a Phred score > 20, variants detected in 20% and less populations, set minimal read depth of seven per allele and finally, filtered to eliminate potential paralogues (SNP rank < 0.4) [53]. To generate the final genetic dataset used for population genetic downstream analyses (*e.g.* SNPs genotype matrix in STRUCTURE, GENPOP or ARLEQUIN format), it was used the component populations.pl implemented in the software STACKS v.1.2 [54] using as input file the resulting filtered VCF above generated by DiscoSnpRad. The filtering parameters in STACKS included that each individual was considered a population in the map file, the minimum number of populations that a locus must be present in to process it was set to 100% (-p = 30) (0% missing data). Other filtering parameters for “-p”, tolerating ∼ 10% and ∼ 5% of missing data in the final dataset (−p = 27 and −p = 29) were also explored to evaluate their statistical power to discriminate samples through PCA analysis, as assessed by the proportion of explained variance, according Martínez et al. (2022) [47]. Likewise, the minimum percentage of individuals in a population required to process a locus for that population was set to 100% (−r = 1), a minor allele frequency (MAF) of 0.01 and retained only one SNP per locus. All other parameters for the analysis were kept as default settings. Finally, we inferred linkage disequilibrium (LD) among SNPs aiming to remove redundant ones from the dataset. To accomplish this, an analysis was performed in the ‘poppr’ R package [54] to evaluate for the existence of linked loci in the SNPs matrix. It was done with the ia() and the pair.ia() functions, which, respectively, calculate the index of association (IA) over all loci in the dataset and calculate the index of association for all pairs of loci in the dataset. A squared correlation between allelic values at two loci - R2 (rbarD) > 0.2 in both the overall loci calculations and ≥10% of pairwise loci comparisons was the criterion for removing linked loci from the matrix [53,55]

In order to obtain the final genomic sequences data matrix (.alleles or .nexus formats) to run analyses based on the coalescence (gene flow, demography and phylogeography), the software PYRAD [56] was used. Two independent runs were done on pyRAD. First, to generate the final genetic dataset used for phylogeographic reconstruction analyses-based, pyRAD was run with the following parameters: Mindepth=10, Wclust=0.88, Datatype=ddrad, MinCov=12, MaxSH=15, PhredScore=20, maxIndels=3,99 and min length trimmed reads=270 bp. These parameters correspond, respectively, to a minimum coverage of 10x for a cluster construction, a clustering threshold of 88% similarity, a requirement that at least 20 samples have data to retain a locus, a maximum number of individuals being heterozygous for a particular base, a minimum quality per base of 20 in the reads based on the Phred Score scale, a maximum number of 3 indels within each cluster construction and up to 99 across clusters, and finally a minimum read length threshold of 270 base pairs for loci to be retained in the final dataset. MaxSH and MaxH serve as filters preventing potential paralogs being included in a cluster. In the second run, to generate the final dataset used for gene flow and historical demography analysis, we set up the same parameters, except for maxIndels, that was set to “0,0”, indicating a requirement of total exclusion of indels within each cluster construction and across clusters to generate the final loci dataset. Tha RADseq raw demultiplexed dataset has been submitted to NCBI as Short Read Archive: accession number PRJNA1308113.

Microsatellites were checked for null alleles using the software Microchecker [57]. The first analysis conducted using the complete data set (SNPs and microsatellites) was an analysis of genetic homogeneity among samples collected in different localities within the Amazon and Orinoco basins, using an Analysis of Molecular Variance (AMOVA) and *F_ST_*[58] (S2 Table). This was done to verify the hypothesis of panmixia within the basins, a fundamental prior to being able to run analyses based on the coalescence including Ima2 and BEAST/Relaxed Random Walk and BEAST/Bayesian Skyline Plot. Since the hypothesis of genetic homogeneity was not rejected (data not shown), samples from Guiana (Guay), Puerto Carreño (PCar), Puerto López (PLop) and Guaviare (Guv), were grouped as Orinoco basin; samples from Boa Vista (BV), Manaus (Mana), Tefé (Tef) and La Pedrera (LPe), were grouped as Amazon basin.

The software STRUCTURE v.2.3.4 [59] and BAPS v.6.0 (*Bayesian Analysis of Genetics Population Structure*) [60] were used to infer patterns of population structure for both nuclear (SNPs and microsatellites) and mitochondrial markers (COI+D-loop), respectively. The analysis implemented in Structure was used to examine the biological groups existing within our sample. Parameters included 1,000,000 Markov Chain Monte Carlo (MCMC) steps and 100,000 iterations discarded as burn-in, an admixture model and correlated frequencies. We explored the possibility of our sample containing from 1 to 8 biological groups (K). Each analysis was repeated 10 times for each simulated K value. The α values and profile of posterior probabilities were examined to assess the convergence between independent runs. STRUCTURE Harvester [61] was used to extract the Q values from each of the 10 independent runs for each K value, and then, summarized in the program CLUMPP v.1.1.2 [62]. Finally, DISTRUCT v.1.1 [63] was used to plot and visualize the Q-matrix obtained, displaying the ancestry probability of each individual in each predefined population. The “Evanno” method [64] was used to infer the most likely number of biological groups (K). In addition, the method of Puechmaille (2016) [65] was used (web server Structure Selector, https://lmme.ac.cn/StructureSelector/) to improve inferences of the number of population clusters offering a better complement that enhances the detection of possible hierarchical clusters (including problems with uneven sampling), not visible with the Evanno. Puechmaille presents strategies for subsampling and performs new analysis methods [the median of medians (MedMed), the maximum of medians (MaxMed), the median of means (MedMean), and the maximum of means (MaxMean)], correcting the weakness of Evanno.

In BAPS, we inferred population structure based on nucleotide frequencies of the mitochondrial region (COI+D-loop) to assignment of individuals. The method provides the posterior probabilities for different numbers of clusters of individuals (K). We performed individual level mixture analysis for multiple defined clusters (K = 1-8 clusters), with 10 independent runs for each K value. The K with the highest posterior probability was selected as the most likely population partition.

To determine if the groupings observed in STRUCTURE or BAPS for each basin were due to recent gene flow or it was the result of ancestral polymorphism retention, we implemented the isolation-with-migration approach in Ima2 [66], to estimate migration rates for *B. rousseauxii* between the basins and its significance. For this analysis, microsatellite data were excluded, considering that: a) the limitations and tendency to violations of IM programs of various assumptions for microsatellite mutation models [67], and b) the possible excess of homoplasy, especially when the divergence/coalescence time between comparing populations is high [68], as in this case. Effective population sizes (*Ne*) were also estimated for the Amazon, Orinoco and for the ancestral basins, depending on the divergence time. First, we ran 50 chains with dynamic heating (-ha 0.99 -hb 0.45) and uniform priors (-q 100 -m 100 -t 100), generating a total of 10,000,000 topologies from which were collected 1%. After an initial burn-in period of 2 million, when parameter estimates stabilized, the topologies were collected. To verify convergence of parameter estimates, we examine the effective sample size (ESS) of the parameters by analyzing trendline plots and comparing parameter estimates based on the first 50% and second 50% of the MCMC run. IMa2 tested the probability of gene flow against the full isolation model using the Log likelihood ratio (LLR) test.

To make the conversions of each scaled parameter (calculated with all markers) into biological data, a conservative generation time of 5 years was assumed for *B. rousseauxii*, consistent with life-history data reported across multiple Neotropical populations. Age at first sexual maturity ranges from approximately 2.2 years in the Madeira River basin to 3–4 years in the Caquetá River (Colombia) and the Peruvian Amazon [69–71]. Given a natural mortality rate of M ≈ 0.50 year⁻¹[69] and applying standard cohort-based generation time estimation (T = α + 1/Z), the expected generation time falls within the range of 5–6 years, which is further supported by the observation that individuals older than 5 years are exclusively found in the headwater reproductive zones of the western Amazon [70], indicating that effective reproductive contribution begins around this age. A longevity of 11–13 years (Agudelo et al. 2013) [71] is also consistent with a generation time of approximately 5 years in a species subject to high fishing mortality [72], given that elevated total mortality rates (Z = M + F) reduce the mean age of reproduction relative to maximum lifespan.

We also assumed a conservative estimate of substitution rate of 0.68x10^-8^ mutations per site per year for the mitochondrial region derived from the widely accepted mtDNA substitution rate for poikilotherm vertebrates in Martin & Palumbi (1993) [73]. For ddRADseqs, we assumed a substitution rate of 1.0x10^-9^ mutations per site per year [74].

The mitochondrial substitution rate adopted here (0.68×10⁻⁸ mutations per site per year) represents a conservative and previously validated estimate for Neotropical freshwater teleosts. This rate was empirically estimated by Hrbek & Larson (1999) [75] from mitochondrial sequence data in Neotropical killifishes and has subsequently been applied in phylogeographic studies within the Orinoco-Amazon biogeographic framework, recovering divergence times consistent with independently documented Plio-Pleistocene connectivity events between these basins [15]. Furthermore, recent large-scale empirical analyses across teleost fishes recovered average mitochondrial substitution rates remarkably similar to the value assumed here [76], providing additional support for the continued applicability of this calibration in freshwater teleosts.

On the other hand, the nuclear substitution rate adopted for ddRADseq loci (1.0×10⁻⁹ mutations per site per year) was selected as a conservative and also was previously validated estimate for Neotropical teleost fishes. This rate has been explicitly applied in previous phylogeographic and divergence-time studies of Siluriformes and Characiformes from South America, yielding chronologies consistent with independently documented geological and hydrographic events [47,77,78]. Furthermore, recent empirical estimates of vertebrate nuclear mutation rates based on pedigree and germline data place ray-finned fishes within a comparable range [79–81], supporting the use of this value as a conservative calibration for nuclear molecular clock analyses in tropical freshwater teleosts.

For the spatio-temporal reconstruction of the ancestral origin and dispersion routes of *B. rousseauxii* between the Amazon and Orinoco basins, a phylogeographic analysis using a Bayesian approach and the *Relaxed Random Walk* model, was run on BEAST v.1.8 [82], in the CIPRES platform [83]. This method infers an evolutionary history into a continuum landscape based on the DNA sequences and the geographic coordinates, generating a genealogy and estimating the ancestral location of the internal nodes, considering uncertainties of the topology [84]. Two of these analyses were conducted, one for mitochondrial sequences (COI + D-loop) and another for nuclear data (ddRADseqs).

For the mitochondrial markers, the substitution model selected was HKY+GAMMA 4 "estimated", and GTR+GAMMA 4 "estimated" for the nuclear markers; both were previously obtained from JModelTest [85]. For the nuclear markers: (a) unphased loci were used, since allele phasing has minimal impact on phylogenetic reconstruction from targeted nuclear gene sequences such as those derived from ddRADseq [86], although ambiguities were retained; and (b) loci were linked for the site model and clock model, but not for the trees among loci. For both analyses, a monophyly prior was imposed on both population clusters based on an initial phylogenetic analysis. The clock model used was an uncorrelated lognormal relaxed clock (non-estimated). The ucld.mean prior was set to a Normal distribution (mean = 0.029, standard deviation = 0.0023), following Sánchez-Bernal et al. (2023) [78], who applied the identical RRW framework to resolve Amazon-Orinoco connectivity in the Neotropical fish *Paracheirodon axelrodi* using the same substitution rate values for poikilotherm vertebrates. A Normal prior was assigned to the population size parameter. All remaining priors — including the spatial diffusion rate precision parameters of the RRW model — were kept at BEAST default settings (Gamma-distributed priors), as recommended by Lemey et al. (2010) [84]. Two alternative coalescent tree priors were tested: Constant Size and Bayesian Skyline.

To select the most likely tree prior scenario, we compared the two alternative models (Bayesian Skyline and Constant Size) using the log marginal likelihood estimation (MLE) obtained by stepping- stone (SS) sampling [87], as implemented in BEAST v.1.8.2. The Bayes Factor was calculated following Kass & Raftery (1995) [88] as Log Bayes Factor0: LBF = 2 × [ln(MLE Bayesian Skyline) − ln(MLE Constant Size)] where a positive LBF value indicates support in favor of the Bayesian Skyline model (Model 1), and a negative value indicates support in favor of the Constant Size model (Model 0). Interpretation of support strength followed the scale of Kass & Raftery (1995): LBF > 10 indicates decisive evidence in favor of the better-supported model.

The continuous RRW model was preferred over discrete phylogeographic approaches (e.g., Bayesian Stochastic Search Variable Selection, BSSVS) because our objective was to reconstruct the macrobiogeographic pattern of colonization — origin, direction, and timing — across the South American subcontinent rather than to quantify transition rates among predefined discrete locations.

RRW in continuous space does not require a priori definition of discrete states, avoids the sensitivity of discrete models to sampling scheme and number of locations [89], and has been validated for this exact biogeographic question in Neotropical freshwater taxa using the same basin-scale spatial extent [47,78].

For the temporal calibration of the phylogenetic tree resulting from each analysis, a substitution rate of 0.68x10^-8^ mutations per site per year for the mitochondrial region, and 1.0x10^-9^ for the ddRADseqs, chosen as explained above. Three independent runs were done, with a chain length of 500 million for the mitochondrial markers and of 250 million for the ddRADseqs, sampled every 100.000 steps. Chain convergence, stationary behavior of the chains and Effective Sample Size (ESS) > 300, were examined in Tracer v1.6.0 and the resulting trees were summarized on TreeAnnotator v1.8.2.

Then, the two independents ran, were combined. To generate spatio-temporal projections of the genealogies in keyhole markup language (kml), the software SpreaD3 v0.9.6 [90] was used with default parameters, and results were accessed and visualized on Google Maps for verification purposes.

We used the program BEAST v.1.8.2 [91] to investigate patterns of changes in effective population size throughout the coalescent time of *B. rousseauxii* in each basin, using the Coalescent: Extended Bayesian Skyline Plot approach (Model 1) and evaluated against the alternative null model Coalescent: Constant Size (Model 0) through log marginal likelihood estimation (MLE) and Bayes Factor comparison (LBF), as described above for the RRW analyses, using Path Sampling (PS), as recommended by Baele et al. (2012). All analyses were run for 50 million generations with a burn-in of 10% initial samples, sampling every 10,000 topologies. We used the HKY+GAMMA model of molecular evolution for mitochondrial (D-loop), and a lognormal relaxed molecular clock. Priors were adjusted based on preliminary analyses. Final analyses were repeated two times to assure convergence of estimated parameters, and the independent runs were combined. Chain convergence, stationary behavior of the chains and Effective Sample Size (ESS) > 300, were examined in Tracer v1.6.0 and the resulting trees were summarized on TreeAnnotator v1.8.2.

The absolute time scale of the Extended Bayesian skyline plot was calculated using a substitution rate for mitochondrial DNA explained and justified above. To evaluate whether demographic change provided a better explanation of the observed genetic data than long-term demographic stability, model support was assessed using the LBF. Positive LBF values indicate support in favor of the Extended Bayesian Skyline model (Model 1), consistent with historical changes in effective population size through time, whereas negative LBF values indicate support for the Constant Size model (Model 0), consistent with long-term demographic stability [92].

### Criteria for strategic selection of individuals for ddRADseq genotyping and hierarchical analysis design

The ddRADseq dataset (n = 30) was not intended to characterize fine-scale within-basin structure—a role served by the microsatellite dataset (n = 154)—but rather to provide genome resolution for evaluating inter-basin differentiation and genealogical history. Individuals for ddRADseq were selected to include balanced representation from both basins (n = 15 per basin) and to specifically oversample the same individuals from some locality identified by SSR analyses as a site showing inter-basin admixture (nuclear ancestry). This strategic inclusion allowed the SNP/ddRADs data to independently corroborate on the same individuals the SSR-based admixture signal (see S1 Table) and provided IMa2 with samples most likely to reveal evidence of recent gene flow, should such gene flow be present. For IMa2, complete flanking sequences loci were used rather than SNPs, providing genealogical information at the topological level required by the MCMC genealogical sampler implemented in the program [66]; under this multi-locus sequence design, inference accuracy for divergence time and migration rate scales primarily with the number of independent loci rather than with per-population sample size [93,94]. Effective population size (*Ne*) estimates from IMa2 are acknowledged to be sensitive to sample size [95] and are therefore interpreted only as relative values (Amazon:Orinoco ratio), not as absolute quantities.

## Results

For the genomic data, a total of 6.81 million reads were obtained, with an average size of 384 bp. Each individual had on average ∼ 0.2 million reads, representing 21373 variable loci. After extracting SNPs and filtering the VCF, 1738 SNPs (0% missing data), representing an equal number of loci, were retained (S3 Table). This dataset corresponded to panel (A) in S2 Fig, and was selected for downstream analyses as it best explained the variance among samples by PCA relative to the other two missing data thresholds evaluated — (B) 5% missing data (1,804 SNPs) and (C) 10% missing data (1,892 SNPs).

Mean depth per SNP in each individual ranged between 14.5 (sample 3847) and 26.5 (sample 21BfCG). After reviewing of LD, we obtained a R2 (rbarD) of 0.0251631 (R2 << 0.2), suggesting that no LD is occurring between loci in the SNP dataset. A heatmap was generated showing paired comparisons of the R2 (S1 Fig).

A total of 12 microsatellite loci (≤15% missing data) were obtained for 154 individuals, and were used for population structure analyses (S4 Table). No null alleles were detected after checking on Microchecker. On the other hand, for phylogeographic analyses, we extracted 326 ddRADseq loci (>270 pb), totaling 94132 bp per individual. For demographic and geneflow analyses (dataset without indels), we extracted 242 ddRADseq loci (484 alleles >270 pb; without missing data across individuals). Finally, we generated 528 and 610 bp of the COI and D-loop regions, respectively, obtaining a total of 1140 bp.

STRUCTURE suggested the existence of two biological groups (K = 2) structured between basins, both for the microsatellites (Ln Pr X|K = -6236.14) and for SNPs (Ln Pr X|K = -26035.5) (Fig 2a-2b) and confirmed by Puechmaille method by the MedMedK=2,MedMeanK=2, MaxMedK=2, MaxMeanK=2 (S3 Fig). Nevertheless, microsatellites indicated admixture between the populations of the two basins in the Boa Vista locality (upper Branco River, Amazon basin) on the Guiana Shield, showing Amazonian individuals with an Orinoco’s genetic composition varying between 19-44%. Population admixture was also inferred in BAPS analyses (Fig 2C), although the most likely number of mitochondrial groups was estimated at K = 4 (log Pr X|K = -872.653).

**Fig 2.**
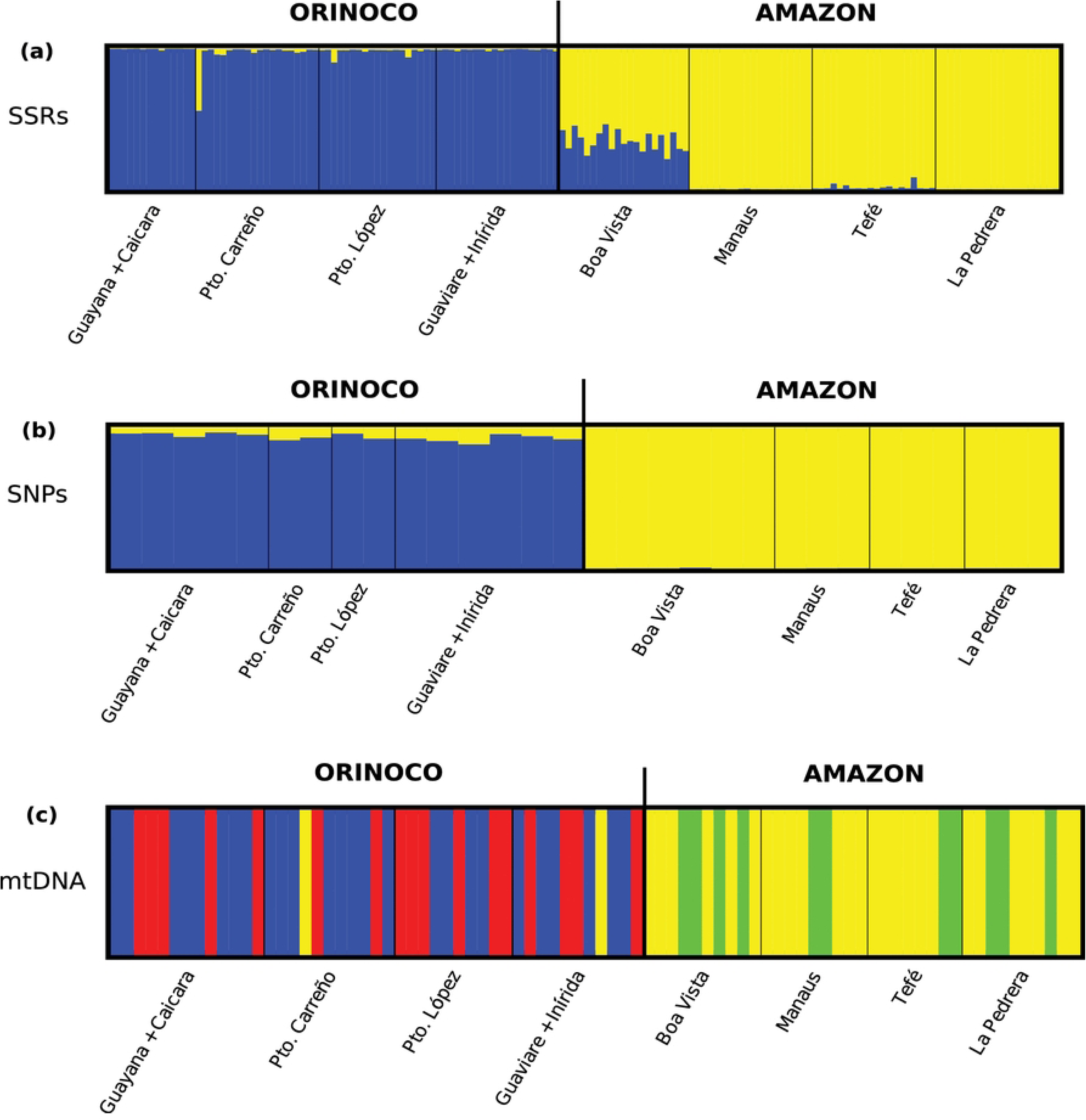
Patterns of genetic structure. Structure analyses of *Brachyplatystoma rousseauxii* samples from the Orinoco [Ciudad Guayana (1), Caicara del Orinoco (2), Puerto Carreño (3), Puerto López (4), Guaviare (5), Inírida (6)] and Amazon basins [Boa Vista (7), Manaos (8), Tefé (9), La Pedrera (10)]. (a) and (b) were inferred by STRUCTURE software, and (c) by BAPS (*Bayesian Analysis of Population Structure*). In (a) and (b), the graphs show the genetic ancestry of each individual, and therefore each bar indicates the probability of an individual belonging to that genetic/biological group either Orinoco or Amazon, represented by blue and yellow, respectively. In (c), the graph shows the more likely cluster arranging of individuals based on the detected genetic mitochondrial groups, which are represented by colors like blue, yellow, red and green.

The Ima2 analyses evidenced that the current gene flow (2Nm) between the two basins is not significant in both directions, showing divergence times of 2.95 and 0.71 Ma, respectively for each marker (Table 1). Both mtDNA and ddRAD markers showed evidence of effective population size expansion in the current Amazon *B. rousseauxii* population relative to the ancestral population.

**Table 1.** Demographic parameters for *Brachyplatystoma rousseauxii* in the Orinoco and Amazon Basins.

| | | $N_e$ (Or)** | $N_e$ (Am)** | $N_e$** (Ancestral) | $2Nm$ (Or → Am) | $2Nm$ (Am → Or) | t (my) |
| --- | --- | --- | --- | --- | --- | --- | --- |
| MLE | mtDNA | 1.981 | 1.829 | 0.485 | 0.179* | 0.195* | 0.706 |
|  | ddRADs | 0.019 | 0.056 | 0.043 | 0.012* | 0.493* | 2.948 |
| Up95% HPD | **mtDNA** | **3.507** | **3.502** | **4.667** | **9.288** | **9.991** | **2.317** |
|  | ddRADs | 0.029 | 0.081 | 0.286 | 0.202 | 0.945 | 3.068 |
| Lo95% HPD | **mtDNA** | **1.172** | **1.027** | **0.000** | **0.000** | **0.000** | **0.442** |
|  | ddRADs | 0.014 | 0.045 | 0.017 | 0.000 | 0.254 | 0.585 |

Maximum-Likelihood Estimates (MLE) and the 95% Highest Posterior Density (HPD) intervals of demographic parameters for the gilded catfish *Brachyplatystoma rousseauxii* from Orinoco (Or) and Amazonas (Am) Basins calculated on Ima2 for sequences data.
\* Non-significant value for the Log-Likelihood Ratio (LLR) test. \*\* Values in millions of individuals.

Conversely in Orinoco, only mtDNA showed expansion, while ddRAD markers showed a current relatively reduced population. Effective population sizes were always greater in the Amazon than in the Orinoco basin based on ddRAD markers estimates, showing a ∼3:1 ratio. By contrast effective population sizes of the mitochondrial genes were approximately equal for the two basins (Table 1).

The Relaxed Random Walk (RRW) phylogenetic reconstruction (Figs 3–4) was explored under two alternative coalescent priors. Model comparison via stepping-stone sampling showed decisive positive support (Kass & Raftery 1995) [88] for the Bayesian Skyline model over the Constant Size model for both datasets: for nuclear markers (ddRADseqs), MLE(Bayesian Skyline) = −7602.88 and MLE(Constant Size) = −7614.11, yielding LBF = 22.465; for mitochondrial markers (COI + D-loop), MLE(Bayesian Skyline) = −3460.12 and MLE(Constant Size) = −3467.36, yielding LBF = 14.470. Both values far exceed the threshold of 10 that Kass & Raftery (1995) define as decisive, consistently favoring the Bayesian Skyline model across both independent marker datasets. The RRW analysis estimated that the most likely ancestral population of *B. rousseauxii* in the Orinoco was located at the Rupununi Portal, in the Guiana Shield (Amazônia), and that the vicariant event that fragmented this population and initiated the colonization of both basins occurred approximately 2.54 Ma (ddRADseqs) or 1.31 Ma (mtDNA). Subsequently, *B. rousseauxii* colonized the Branco River and the Orinoco River delta at 1.90 Ma (ddRADseqs) or 0.6 Ma (mtDNA), expanding rapidly throughout the Orinoco basin.

**Fig 3.**
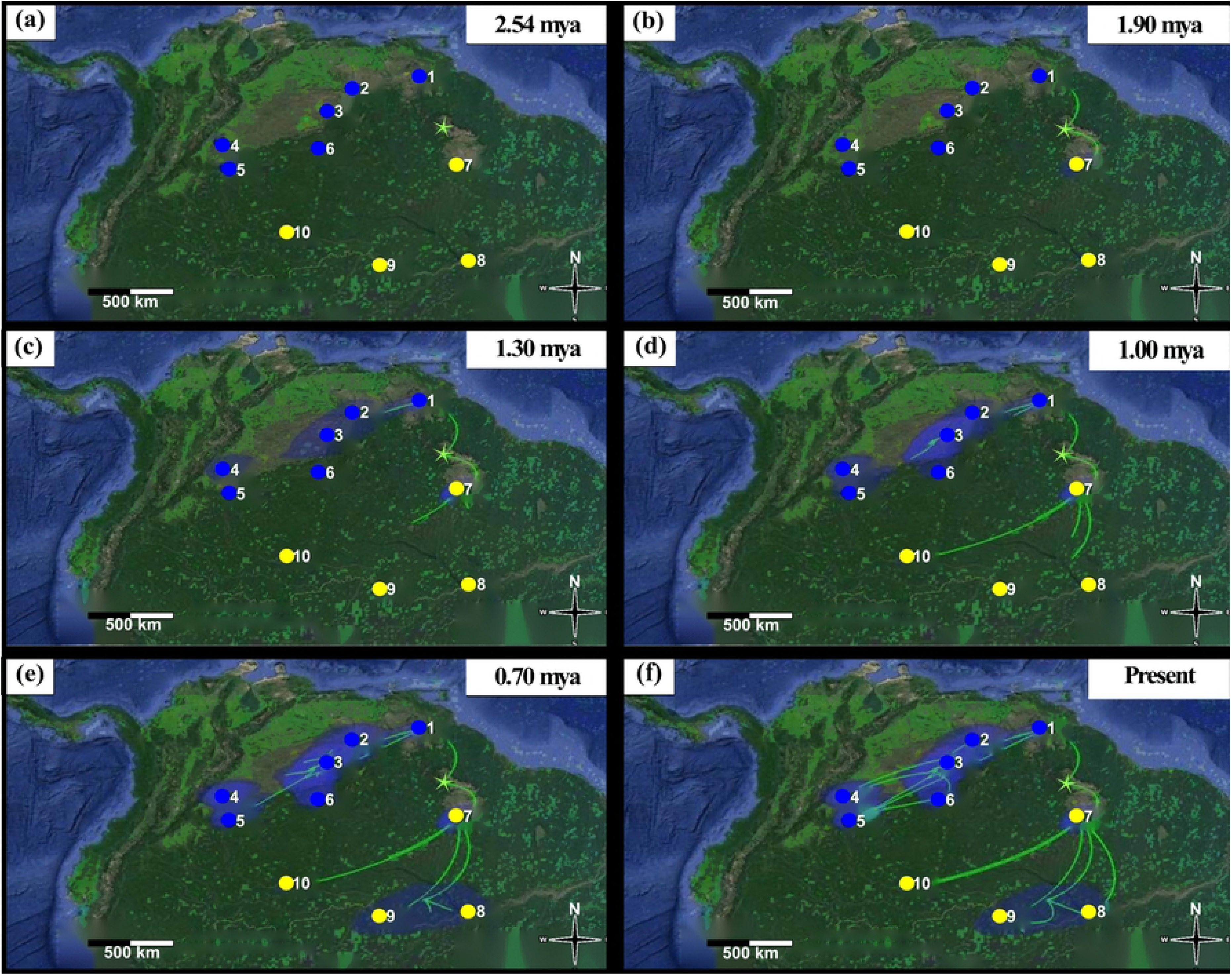
Phylogeographic reconstruction of ancestral area using nuDNA. Coalescent reconstruction analysis using ddRADseqs dataset, based on a relaxed random walk in continuous space and time for *Brachyplatystoma rousseauxii* from Orinoco (blue circles, localities 1–6) and Amazon (yellow circles, localities 7–10) basins, inferred in BEAST and visualized in SPREAD3. Graph is showing the hypothetical spatial patterns and times of colonization for *B. rousseauxii* in each basin, locating the ancestral population from all of them were originated (green star). All the observed colonization times are median heights into 95% credibility intervals, which can be consulted in the supporting information in S4 Fig. Green lines represent the branches of the Maximum Clade Credibility (MCC) tree projected onto geographic space, depicting the inferred dispersal trajectories of *B. rousseauxii* lineages from ancestral to descendant geographic locations through time; the direction of each branch follows the inferred colonization route. Blue polygons represent the 80% Highest Posterior Density (HPD) contours for the inferred geographic location of internal nodes (ancestral areas) at each time slice, reflecting the spatial uncertainty of the estimated ancestral positions. Map and background raster data were obtained exclusively from publicly available datasets licensed under the Creative Commons Attribution 4.0 (CC BY 4.0), including Landsat (http://landsat.visibleearth.nasa.gov/) and the USGS Earth Resources Observation and Science (EROS) Center (http://eros.usgs.gov/#). No copyrighted or proprietary data sources (e.g., Google Maps or Google Earth) were used in the creation of this figure. The map was constructed using these datasets within the R statistical environment (www.r-project.org) and subsequently edited using Inkscape v1.X (https://inkscape.org).

**Fig 4.**
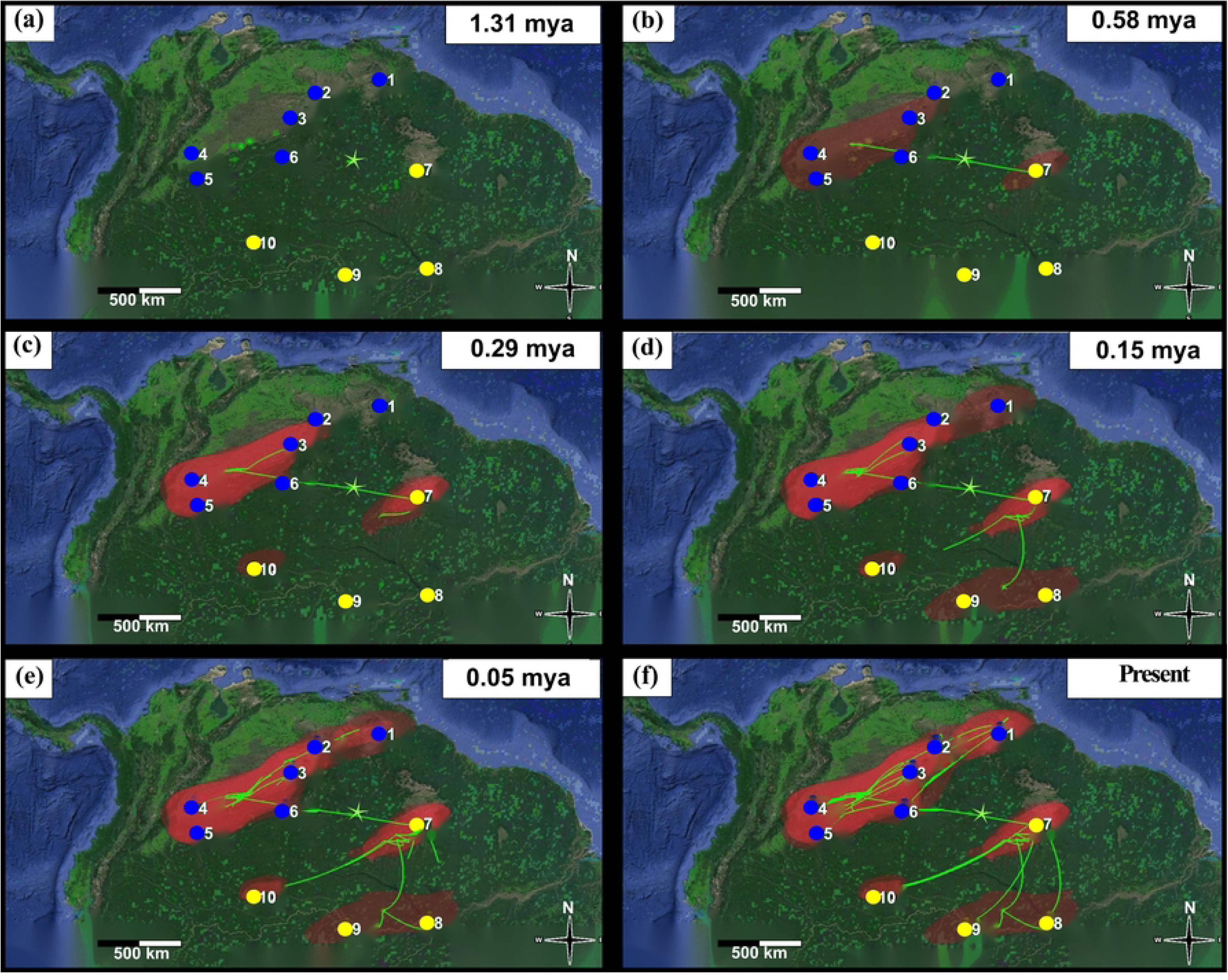
Phylogeographic reconstruction of ancestral area using mtDNA. Coalescent reconstruction analysis using mtDNA concatenated dataset (COI+D-loop), based on a relaxed random walk in continuous space and time for *Brachyplatystoma rousseauxii* from Orinoco (blue circles, localities 1–6) and Amazon (yellow circles, localities 7–10) basins, inferred in BEAST and visualized in SPREAD3. Graph is showing the hypothetical spatial patterns and times of colonization for *B. rousseauxii* in each basin, locating the ancestral population from all of them were originated (green star). All the observed colonization times are median heights into 95% credibility intervals, which can be consulted in the supporting information in S5 Fig. Green lines represent the branches of the Maximum Clade Credibility (MCC) tree projected onto geographic space, depicting the inferred dispersal trajectories of *B. rousseauxii* lineages from ancestral to descendant geographic locations through time; the direction of each branch follows the inferred colonization route. Red polygons represent the 80% Highest Posterior Density (HPD) contours for the inferred geographic location of internal nodes (ancestral areas) at each time slice, reflecting the spatial uncertainty of the estimated ancestral positions. Map and background raster data were obtained exclusively from publicly available datasets licensed under the Creative Commons Attribution 4.0 (CC BY 4.0), including Landsat (http://landsat.visibleearth.nasa.gov/) and the USGS Earth Resources Observation and Science (EROS) Center (http://eros.usgs.gov/#). No copyrighted or proprietary data sources (e.g., Google Maps or Google Earth) were used in the creation of this figure. The map was constructed using these datasets within the R statistical environment (www.r-project.org) and subsequently edited using Inkscape v1.X (https://inkscape.org).

Only much more recently did the species expand from the Branco River to the rest of the Amazon basin, at approximately 1.0 Ma or 0.15 Ma, respectively for each marker.

The Extended Bayesian Skyline Plot (Fig 5) was explored under two alternative coalescent priors and compared via Path Sampling marginal likelihood estimation (Kass & Raftery 1995) [88]. For the Amazon population, MLE(Extended Bayesian Skyline) = −1957.95 and MLE(Constant Size) = −1953.81, yielding LBF = −8.28; negative LBF values indicate support for the Constant Size model (Model 0), consistent with long-term demographic stability. For the Orinoco population, MLE(Extended Bayesian Skyline) = −2072.61 and MLE(Constant Size) = −2077.69, yielding LBF = +10.16, exceeding the decisive threshold of 10 (Kass & Raftery 1995) [88] and thus favoring the EBSP (Model 1), consistent with the hypothesis of historical demographic change. The median *Ne* trajectory observed in the EBSP plot suggests a probable mild demographic expansion in the Orinoco population, though the 95% HPD intervals are wide and the precise magnitude and timing of *Ne* changes carry inherent coalescent uncertainty. ESS > 300 was confirmed for all parameters in Tracer v1.6.0. Extended Bayesian skyline plots showed that the more recent common ancestor for Orinoco *B. rousseauxii* population was detected at ∼1.1 million years before present (YBP), while in the Amazon basin it was detected much more recently at ∼500,000 YBP. Additionally, it was evidenced that the population size of *B. rousseauxii* has remained constant in the Orinoco basin until approximately 250,000 YBP, after which point has been experiencing a moderate demographic growth (Fig 5a). On the other hand, the Amazon basin exhibited a relatively stable demographic history of *B. rousseauxii* through time up to the present (Fig 5b).

**Fig 5.**
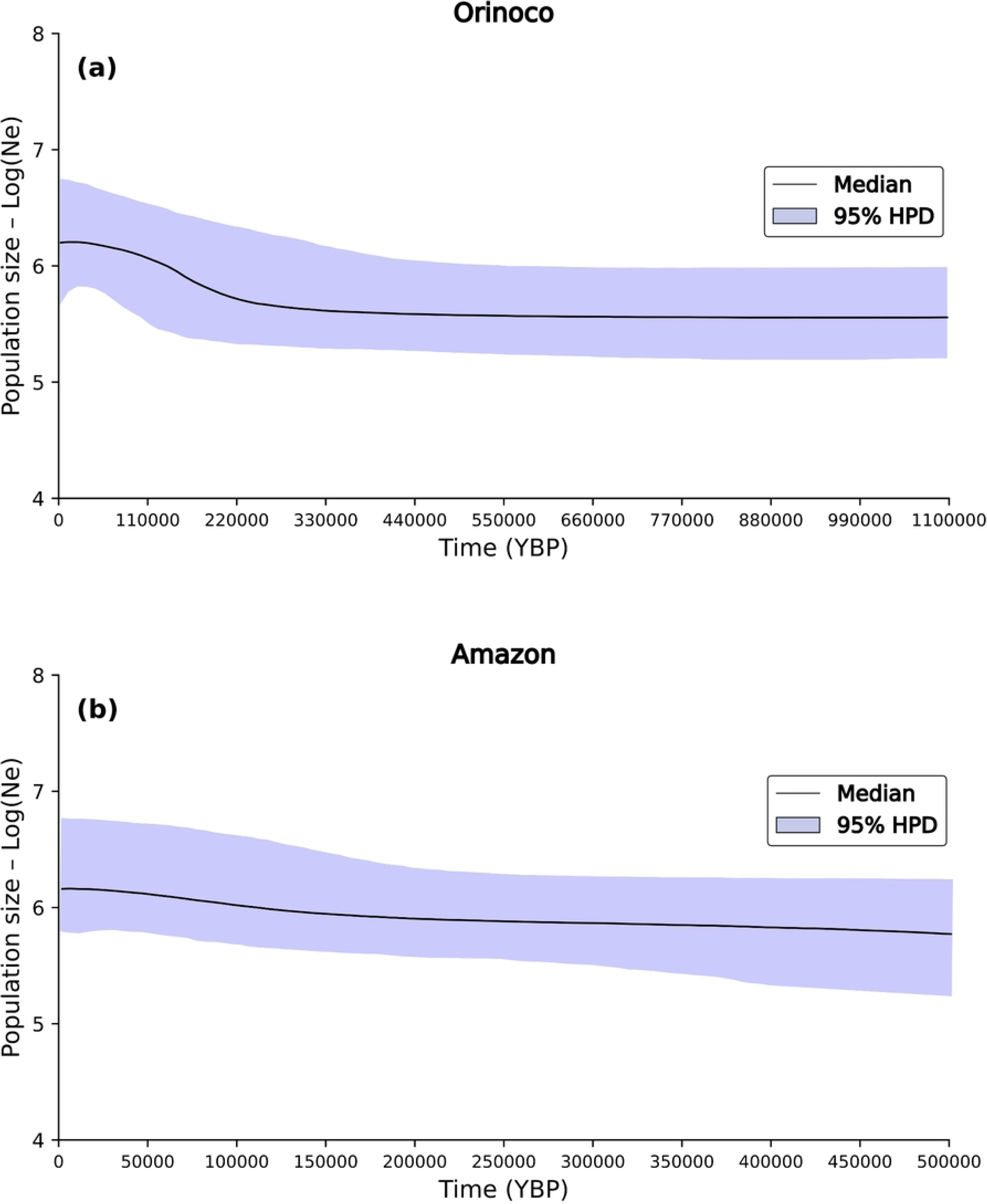
Skyline Plot historical demographic reconstruction. Extended Bayesian skyline plot analysis inferred in BEAST and estimated from Orinoco (a) and Amazon (b) samples using the mitochondrial D-loop region. The thick solid line is the median estimate, and the limit defined by violet space represents the 95% Highest Posterior Density limits. Note: *Ne* is on a log10 scale.

## Discussion

### High structuring and no current connectivity for *B. rousseauxii* between basins

Here, microsatellites, SNPs ddRAD-based, and mtDNA markers provided strong evidence suggesting that the Vaupés Arch and Guiana Shield physical barriers currently act as a strong obstacle for ongoing gene flow and dispersion of *B. rousseauxii* between the Amazonas and Orinoco basins, generating a high current genetic structure and suggesting a vicariant event distributing the genetic diversity between basins.

The variation patterns generated by these barriers have been previously observed in sister species of populations of a particular species inhabiting each basin, for example, sister species of piranhas *Pygocentrus nattereri/P. cariba,* or cichlids*, Uaru amphiacantoides/Uaru fernandezyepezi* and *Biotoecus opercularis*/*B. dicentrarchus.* Also, for populations of *Piaractus brachypomus* or *Serrasalmus rhombeus* [14,19]. The rise of these barriers has been the most likely explanation supporting the distribution and diversity of ichthyofauna found between the Amazon and Orinoco, including endemism, in which one taxon is found in one basin and absent in the other, as it has been found for the lungfish *Lepidosiren paradoxa*, pirarucu *Arapaima gigas* and for cichlids of the genus *Symphysodon* [4]. Today, due to these barriers, of the total combined ichthyofauna recorded in both basins, 61.2% of species are exclusive to the Amazon, 16.6% are exclusive to the Orinoco, and 22.2% are shared between the two basins [14], suggesting that the biogeography of these regions are complex, involving not only vicariance but dispersal across some portals that may limit or not the species distribution, which will depend on its own morphological or ecological capacity to cross the portals [5,11].

Due to its body size, migration capacity and broad distribution range between Orinoco and Amazon basins [28,96], *B. rousseauxii* might be expected to facilitate recent inter-basin gene flow. However, in this study neither the Japurá/Guaviare, Casiquiare, nor Rupununi portal resulted in current connectivity routes between Orinoco and Amazon populations, as shown by IMa2. In fact, the barriers or events causing the highly observed genetic structuring seem to be ancient, appearing strong and persistent since the early (2.94 Ma) or late (0.71 Ma) Pleistocene until the present days.

However, this date does not coincide with the rise of Vaupés Arch, the most commonly used orogenic event in the Miocene to explain Orinoco and Amazon basin separation and vicariant speciation [9,97] thought to have occurred about 10 to 8 Ma, extending its elevation up to early Pliocene [98] about 5.3 Ma. Instead, it coincides with the age for the final stage of the onset of the transcontinental Amazon River, about 3–0.78 Ma [99]. This final stage (Stage 2, ca. 4.5–0 Ma) involved the breaching of the Purús Arch — a Palaeozoic structural high forming the eastern watershed of the sub-Andean foreland — which redirected western Amazonian sub-Andean drainages eastward to the Atlantic and effectively separated them from the northern Orinoco drainage system [99]. Optically stimulated luminescence (OSL) dating of river channel deposits places this large-scale drainage reorganization at approximately 2.6 Ma [99], directly coinciding with the molecular divergence window estimated for *B. rousseauxii*. Concurrent Northern Andean surface uplift, concentrated between ∼8 and ∼4 Ma, progressively isolated the Orinoco, Magdalena, and Amazon drainages from their formerly shared proto-Andean foreland predecessor [87,88].

The onset of Quaternary glacial–interglacial cycles at ∼2.6 Ma further modulated inter-basin connectivity in the Amazon–Orinoco lowlands through repeated cycles of glacial river incision and interglacial floodplain expansion [75,87,88]. That these geological and climatic events could have driven vicariance specifically in *B. rousseauxii* is supported by the demonstrated susceptibility of Pimelodidae to river capture events: Tagliacollo et al. (2015) [100] showed that members of this family consistently bear biogeographical signatures of drainage capture throughout their evolutionary history across South American river systems. Consistent with this, the fossil-calibrated molecular phylogeny of those authors recovered intraspecific divergences within three wide-ranging goliath catfish species sampled across their full Neotropical distributions — *B. rousseauxii* (0.5 Ma), *B. vaillantii* (0.6 Ma), and *B. filamentosum* (3.7–0.2 Ma) — all concentrated in the Plio-Pleistocene window and calibrated independently of any assumed substitution rate, providing corroborating evidence that Plio-Pleistocene river reorganizations structured intraspecific diversity within *Brachyplatystoma* across basins [100]. Interestingly, this Pleistocene split time between basins for *B. rousseauxii* coincides with estimates reported for other fish species in the same biogeographic realm: *Piaractus brachypomus* (2.54 or 0.44 Ma; [19]), actinopterygian species pairs (2.5 Ma; [101]), and species of the genera *Potamorrhaphis* (2.5 Ma; [18]), *Prochilodus* (1.57 Ma; [102]), and *Austrofundulus* (2.95 Ma; [103]), assembled from independent primary studies in a comparative phylogeographic framework by Escobar et al. (2015) [19].

The concordance in Plio-Pleistocene divergence times across these phylogenetically distinct taxa, each derived from independently measured sequence divergences in separate datasets, is consistent with a genuine biological signal of near-simultaneous vicariance driven by the geological events described above [104]. One of the explanations that could support the lack of current connectivity for *B. rousseauxii* between basins may be related to its ecological inability to use black-water channels such as the Negro River-Casiquiare corridor. The species is exclusively distributed throughout the main channels and white-water tributaries of the Amazon and Orinoco basins [12,105,106], with no confirmed records in the Negro River system. The physicochemical characteristics of this river — the largest black- water river in the world, with strong acidity (pH 4.4–4.9) and extremely low ion concentrations [107–109] compared to white-water systems [110] — suggest that this corridor would impose a significant physiological challenge for a large, obligate carnivore with high energetic demands: tolerance to ion- poor, highly acidic conditions requires specific ionoregulatory adaptations — including high-affinity Na⁺ uptake systems and tight regulation of ion efflux — that have been documented only in certain Amazonian fish orders (Characiformes, Cichliformes), but remain uncharacterized in Siluriformes [111,112]. Non-adapted fish exposed to such conditions suffer net losses of essential ions with severe physiological consequences [Gonzalez et al. 2002]. We acknowledge, however, that no experiments or systematic sampling have directly tested *B. rousseauxii* tolerance to black-water conditions, and this hypothesis remains to be empirically verified. The Casiquiare canal further compounds this barrier, exhibiting an extreme physicochemical gradient from pH 6.7 and clear waters at its Orinoco origin to pH 4.8 and black waters at its confluence with the Negro River [11].

The idea that this corridor acts as a selective ecological filter — permeable to some taxa but not others — is supported by comparative biogeographic and genetic evidence. Species permanently distributed along the Negro-Casiquiare, including the river dolphin *Inia geoffrensis*, the turtle *Podocnemis expansa* [113], the piranha *Serrasalmus rhombeus*, all species of *Boulengerella*, the cichlids *Cichla temensis* and *Mesonauta insignis*, and the catfishes *Pimelodus blochii* and *Scorpiodoras heckelii* [11,14], show ecological tolerance to black-water conditions. Genetic studies confirm inter-basin connectivity through this corridor for black-water-adapted species: *Cichla temensis*, *C. monoculus* and *C. orinocensis* [5], *Osteoglossum ferreirai* [114], and *Paracheirodon axelrodi* [78]. By contrast, white- water specialists such as the turtle *Podocnemis unifilis* [115] and the fish *Piaractus brachypomus* [19] show no current gene flow between basins, consistent with the hypothesis that the corridor functions as an effective barrier for species lacking black-water adaptations — a category in which *B. rousseauxii* most plausibly falls.

In this study, we initially hypothesized that the Guaviare (Orinoco) and the Japurá (Amazon), by being white water rivers with headwaters located in the western Amazon (Andean piedmont), right above the region that constituted the old Paleo-Amazonas-Orinoco River [10,116], would become a hypothesis of irrefutable current connectivity portal between basins. This is, that temporal migration could have been expected during high water seasons by extension of the flood plains, as suggested by Lundberg in a personal communication in 2012. However, our results reject this hypothesis, at least for *B. rousseauxii*.

In spite of the panmixia found in this study for *B. rousseauxii* within the Orinoco and Amazon, which is characteristic for species of this genus [117,118], this species could not have the ability to move in the headwaters of the Japurá (Amazon) and Guaviare (Orinoco) to establish a connection between those rivers, probably by the difficulties in locomotion due to their big size, mainly in headwater systems characterized by small, shallow rivers [119]. Also, it is possible that waterfalls located at Serranía de Chiribiquete (Japurá river, Colombian side) are preventing upstream migration to the headwaters of the Guaviare River. In fact, waterfalls are found along the Japurá River: Araracuara (500 km from the border with Brazil), and then the waterfalls of Angosturas (a further 50 km upstream). Also, there are rapids such as the Yarí, La Sardina and Córdoba [120]. Likewise, along the length of the Japurá river it gets reduce from 1000 m to 120 m, achieving vertical falls of more than 100 m in height [121]. Carvajal et al. (2014) [122] showed that *B. rousseauxii* has restrictions to gene flow between the Bolivian Amazon sub-basin (upstream Madeira River) and at the Solimões river (downstream Madeira River), explaining such patterns by the presence of more than 18 rapids between those areas [123].

Also, even without current connectivity, we expected to have at least some evidence suggesting the Japurá/Guaviare as being the last point of contact in a recent past between both basins. However, through the STRUCTURE analysis we rejected this hypothesis, suggesting that the more likely last point of contact between the two basins was not located in the western Amazon but probably in the Guiana Shield east Amazon. This is much more distant than what one day the Paleo-Orinoco-Amazonas River was, confirming the hypothesis by Lovejoy & Araujo (2000) [18] about the ancient Rupununi Portal.

Although it was a hypothesis initially created to explain the colonization of marine fish species to the Amazon, it could also have provided a connection for freshwater species between these basins, as suggested by the current evidence of hydrological connection between the Guyana Shield and the Amazon basin via the Rupununi savannas and wetlands [24,124].

We found nuclear evidence supporting the existence of that Portal, also proposed by Hubert & Renno (2006) [23]. Even with the absence of current connectivity between basins, the Amazon population of Boa Vista in the Upper Rio Branco, located on the Guianese shield, showed some individuals who clearly retain traces of the nuclear genome of the Orinoco (19-44%), suggesting a historical connection between basins via Rupununi Portal on the Guiana Shield for *B. rousseauxii*. No other Amazonian location other than Boa Vista showed this blunt historical sign of genomic sharing with Orinoco, suggesting that the Upper Branco River was perhaps the last contact area or last stepping point for individuals and the exchange of alleles between populations of the species. Similar results were evidenced in ancestry analysis for pirapitinga *Piaractus brachypomus*, also using microsatellites [19], finding a genome sharing with Orinoco only in individuals from Boa Vista locality varying from 5-34%.

Lasso et al. (1990) [125] showed ichthyological evidence that the Cuyuni River (Guiana Shield) formed part of the Caroni-Orinoco River drainage, captured by the Mazaruni River that actually drains into the Essequibo River. Today, the Essequibo is connected via the flooded savannahs of the Rupununi River to the Branco River which drains into the lower Negro River, direct affluent of the Amazon River. This current hydrological connection (Essequibo-Rupununi-Branco River) allows for ichthyofaunal exchange between these riverine systems [18,25–27].

### Ancestral polymorphism, not contemporary gene flow, explains the Boa Vista admixture signal

The admixture detected in Boa Vista individuals by microsatellite STRUCTURE analysis (19– 44% Orinoco nuclear ancestry) does not contradict the IMa2 finding of non-significant contemporary gene flow. This pattern is well explained by a phenomenon called retention of ancestral polymorphism: shared genetic variation inherited from a common ancestor can persist across populations long after divergence, generating detectable coancestry signals even in the complete absence of ongoing exchange [126,127]. When considered at the level of individual loci, this process is also referred to as incomplete lineage sorting (ILS; [128]) and is particularly pronounced in populations with large ancestral *Ne* or a history of secondary contact followed by renewed isolation— both conditions are likely satisfied in the *B. rousseauxii* metapopulation, as indicated by our results. Microsatellites are especially susceptible to generating such misleading signals, as their high mutation rate and multi-allelic nature make STRUCTURE bar plots prone to reflect deep shared ancestry rather than recent mixing [67,68,129].

Critically, when some of the same genetically admixed Boa Vista individuals were genotyped at genome-wide SNP/ddRADseq resolution (>1,700 SNPs), STRUCTURE assigned them unambiguously to the Amazon cluster, and IMa2 confirmed isolation as the best-supported model (LLR test). The coancestry signal at Boa Vista therefore could be reflecting retained ancestral polymorphism from a Pleistocene secondary contact through the Rupununi Portal, most parsimoniously interpreted as the genomic footprint of connectivity that preceded the final Branco-Essequibo disconnection and subsequent capture of the upper Branco River by the Negro River (∼0.25–1.0 Ma) [130]—a timeframe fully consistent with the Late Pliocene to mid-Pleistocene divergence estimated by both IMa2 (0.71– 2.95 Ma) and the RRW analyses (1.31–2.54 Ma). The possibility of human-mediated inter-basin translocation was also considered and excluded: unlike omnivorous, small-bodied species such as *Piaractus brachypomus* and *Colossoma macropomum* for which established aquaculture technology enables deliberate stocking programs [131]. To date, no evidence has been found in either the scientific or gray literature indicating that *B. rousseauxii* has been successfully reproduced in captivity, and no stock enhancement or restocking programs have been reported anywhere across its distribution range. This likely reflects the biological and logistical constraints imposed by its exceptionally large body size, difficulties in captive maintenance, slow growth, long generation time, and obligate long-distance reproductive migration.

### Mito-nuclear temporal discordance: a biological signal, not a methodological artifact

The divergence times inferred from concatenated mtDNA (COI + D-loop) were systematically younger than those obtained from multi-locus ddRADseq nuclear markers across all coalescent analyses performed in this study. The ancestral population split was dated at ∼1.31 Ma by mtDNA versus ∼2.54 Ma by ddRADseq, while colonization of the Branco and Orinoco rivers is estimated at ∼0.6 Ma versus ∼1.90 Ma, respectively; IMa2 analyses yield a parallel pattern (0.71 Ma versus 2.95 Ma). Crucially, both marker systems recover an identical biogeographic scenario—an ancestral population on the Guiana Shield, earlier and broader colonization of the Orinoco basin, and a more recent entry into the Amazon through the captured Branco River—differing only in the absolute timing of these events by approximately two- to threefold. Such mito-nuclear discordance in divergence time estimates is neither unexpected nor exceptional; it reflects well-documented differences in the evolutionary properties of these two genomes that are amplified when historical secondary contact has occurred.

Although additional and a more in-depth investigation is required, the most fundamental explanation for the consistently younger mtDNA-based divergence estimates likely resides in differences in effective population size (*Ne*). Mitochondrial DNA, by virtue of its uniparental inheritance and haploid nature, has an *Ne* approximately four times smaller than that of biparentally inherited nuclear loci [132,133]. A smaller *Ne* accelerates coalescence, causing the mitochondrial gene tree to reach its common ancestor more rapidly and thus producing apparently more recent divergence times relative to the nuclear genome. This effect is compounded by substitution saturation: mitochondrial markers accumulate substitutions at elevated rates—particularly at third codon positions—which can cause underestimation of deeper divergence times and overestimation of more recent ones, a pattern now empirically documented across multiple tetrapod orders [134]. In freshwater fishes, comparative analyses have similarly demonstrated that mtDNA and nuclear sequence data reveal different demographic histories at different temporal scales precisely because of these *Ne* and rate differences [135].

A second, biologically compelling explanation for the discordance could be the historical mitochondrial introgression through the Rupununi Portal. In their landmark review of 126 animal systems with well-documented mito-nuclear biogeographic discordance, Toews & Brelsford (2012) [133] showed that this pattern most frequently arises after allopatric isolation followed by secondary contact, with the mitochondrial genome disproportionately captured by the nuclear background of an adjacent lineage—an event that effectively resets the mitochondrial coalescence clock to the time of contact rather than the time of original vicariance. In *B. rousseauxii*, our nuclear data provide strong evidence that such a contact event likely occurred, particularly given the absence of contemporary gene flow inferred from IMa2: individuals from Boa Vista (upper Branco River, Amazon basin), located at the gateway of the Rupununi Portal on the Guiana Shield, carry 19–44% Orinoco nuclear ancestry, while no other Amazonian locality shows this signal. If mitochondrial haplotypes were also exchanged during this episode—even partially— the resulting mtDNA coalescence would reflect the more recent contact between basins event rather than the deeper vicariant split, thereby producing the observed pattern of both reduced divergence times and signals of mitochondrial haplotypic proximity as observed in BAPS. This mechanism has been explicitly documented in freshwater teleosts: Perea et al. (2016) [136] demonstrated in the genus *Squalius* (Actinopterygii: Cyprinidae) that ancient mitochondrial capture during Pleistocene secondary contact caused severe mito-nuclear discordance, with multi-locus nuclear data accurately reflecting species boundaries while mtDNA patterns were driven by the direction and timing of historical gene flow.

Importantly, the observed discrepancy cannot be attributed to poorly chosen or unvalidated substitution rates. The mitochondrial rate assumed here (0.68×10⁻⁸ mutations per site per year) was empirically estimated by Hrbek & Larson (1999) [75] from mitochondrial data in Neotropical killifishes (Rivulidae) and was subsequently applied by Escobar et al. (2015) [19] to reconstruct the divergence history of both *Austrofundulus* (Cyprinodontiformes) and *Piaractus brachypomus* (Characiformes) between the Orinoco and Amazon basins—recovering splits of ∼2.95 Ma and ∼2.54 Ma, respectively, fully concordant with known Plio-Pleistocene inter-basin connectivity events in the same biogeographic context as the present study. This rate is further supported by the recent large-scale empirical estimation of mitochondrial substitution rates across 139 teleost species [76], which recovered a mean rate of 0.79×10⁻⁸ substitutions per site per year, strikingly close to our assumed value. Similarly, the nuclear rate (1.0×10⁻⁹ mutations per site per year) has been applied in two independent studies carried out in the Neotropical framework: Martínez et al. (2022) [47] and Ibarra et al. (2026) [77] used it for the nuclear marker rag2 and ddRADs in Pimelodidae catfishes from the Magdalena-Cauca basin, recovering species-level divergences of 0.66–3.32 Ma and temporal demographic origins (∼4.5 Ma) consistent with the known geomorphological dynamics of the inter-Andean valleys; and Sánchez-Bernal et al. (2023) [78] applied it to the nuclear marker MYH6 in *Paracheirodon axelrodi* across the Orinoco and Negro- Amazon basins, obtaining a biogeographic history fully concordant with the formation of the Vaupés Arch and Pleistocene evolution of the Casiquiare Canal and Negro River.

Likewise, germline mutation rate surveys across 68 vertebrate species [79] independently place the average annual nuclear substitution rate for ray-finned fishes at 1–3×10⁻⁹ mutations per site per year given generation times typical for tropical siluriform fishes (2–5 years), bracketing our assumed rate as a well-justified conservative lower bound. The fact that both rates are independently validated in the Neotropical [Andean or Orinoco-Amazon] biogeographic context and produce geologically coherent chronologies when applied to the appropriate marker systems demonstrates that the two- to threefold discrepancy observed here could not be a methodological artifact of rate misspecification, but a likely genuine biological signal of mito-nuclear discordance driven by the differences probably in *Ne*, substitution saturation dynamics, or historical mitochondrial “introgression” (secondary contact) discussed above.

For all these reasons, we consider the ddRADseq nuclear timeline to provide the more reliable chronological framework for the historical biogeography of *B. rousseauxii*. It is based on thousands of independent loci distributed across the genome, buffering against the influence of selection, introgression, or incomplete lineage sorting at any individual marker; and the nuclear dates align closely with independent geological evidence for the capture of the upper Branco River by the Negro River, estimated at ∼0.25–1.0 Ma [130,137], consistent with the ddRAD-based colonization time of the Amazon (∼1.0 Ma) and the detection of the most recent common ancestor within the Amazon basin (∼460,000 YBP by BSP).

The mtDNA could be estimating a remaining valuable as a conservative lower bound on divergence times and as a record of the timing of mitochondrial introgression during late Pliocene– Pleistocene connectivity events through the Rupununi Portal. Indeed, it is also worth noting that the Amazon River itself did not arise as a single instantaneous event. Age estimates for its transcontinental capture span from ∼12.6 Ma to 0.019 Ma [88], reflecting a prolonged history of successive drainage reorganizations across the Neogene and Quaternary. Similar complex histories likely characterize other major Neotropical river systems, suggesting that the distributional patterns of their ichthyofaunas — including that of *B. rousseauxii* — are the product of an equally layered and temporally extended biogeographic process.

### Guiana shield probably acted as an ancient portal and putative ancestral area for *B. rousseauxii*

All important historical biogeographic findings for *B. rousseauxii*, occurred from the very Late Pliocene to mid-late Pleistocene, from 2.95 up to 0.15 Ma. Along this time, the ancestral population of *B. rousseauxii* localized probably on the Guiana Shield, experienced a vicariant event fragmenting it thereby initiating the colonization of both basins, which took place from the highlands of Guiana Shield (Rupununi Portal) to the lowlands of the Orinoco and Amazon basins.

One of the stronger hypotheses to explain the observed vicariant pattern involves the geomorphological dynamism of the lowland margins surrounding the Guiana Shield. Although the Shield itself is a geologically stable Precambrian craton, its peripheral lowlands are subject to ongoing processes of differential headward erosion, sediment deposition, and landscape-driven changes in drainage gradients [98,119,138], all of which can redirect headwater streams and trigger river capture events without requiring any recent tectonic activity. Such stream capture events produce simultaneous vicariant and geodispersal effects — isolating the headwaters of one basin while connecting them to an adjacent one [8,100]; a very studied mechanism for explaining the diversification patterns in Neotropical catfishes [100]. Fouquet et al. (2012) [139] found that the genetic substructure observed in aquatic frogs throughout the Guiana Shield suggests further early Pleistocene (c. 2.6 Ma) fragmentation and a very important role for rivers disturbing/rearranging, supporting fragmentation of moist tropical forest and aquatic environments in the Guiana Shield during this period.

Although a clear highland-to-lowland pattern of colonization was observed for *B. rousseauxii*, the important question is: Why does it begin at 2.54 Ma? A hypothesis explaining this fact is the complete establishing and connectivity of the main Rivers systems in the Neotropics lowlands, about 2.5 Ma [140]. This is proposed to have occurred at the start of the Plio-Pleistocene glacial climate cycle during the lowest sea level, with concomitants geomorphological changes that continued until the Holocene [130,137,140,141]. The final establishment of the main Neotropical River basins in this epoch, with simultaneous decrease of sea levels, could have promoted a more rapid lowland colonization, where possibly new habitats and resources were available. An example of this pattern occurred in the freshwater fish species puffer *Colomesus asellus*, which experienced range expansions and colonization of the available freshwater habitats throughout Orinoco system and in the proto-Amazon. In this last river system, *C. asellus* followed a clear west-east colonization pattern until the late Miocene after which the proto-Amazon River breached the Purus Arch. At that time, approximately 2.5 Ma, the modern Amazon River established with a west-to-east transcontinental flow, and puffers subsequently colonized downstream, following the freshwaters environments now available [142]. Although the continuous RRW model does not capture the fine-scale topological constraints of river networks, future work employing discrete phylogeographic approaches — such as Bayesian Stochastic Search Variable Selection (BSSVS; [89]) — could resolve specific dispersal routes among individual tributaries and quantify asymmetric transition rates between drainage units, providing a complementary and higher- resolution view of the macrobiogeographic patterns described here.

### A too late colonization of Amazonas basin by *B. rousseauxii*

In this study, it was clear that the Orinoco was first colonized and populated by *B. rousseauxii* from the ancestral uplands of the Guiana Shield. We observed strong differences, about 300,000 – 150,000 years, in the colonization/establishment time of *B. rousseauxii* throughout the lowlands of Orinoco Vs. lowlands of Amazon basin. This pattern is corroborated by BSP results, which showed that the more recent common ancestor for *B. rousseauxii* population appears in Orinoco approximately 500,000 years before than in the Amazon.

The explanation may have been associated with the late establishment of an aquatic connection between the uplands of Guiana Shield and the lowland Amazon basin. Until about one million years ago (1 Ma), the upper Branco River drained into the Essequibo River [143], and only in the mid-late Pleistocene, about 0.25–0.1 Ma [130], it was captured by the Negro River to become part of the Amazon basin. Both the period of colonization of *B. rousseauxii* to the Amazon system in the mid-late Pleistocene (1 or 0.15 Ma) and the detection of its most recent common ancestor within the Amazon basin (0.5 Ma) coincide with the transition period between the Branco–Essequibo disconnection and the subsequent final capture of the Branco River by the Negro River and the establishment of its current course, estimated at 1–0.1 Ma [130,143], a phenomenon that could have facilitated the arrival of this species in the Solimões, and from there, to the rest of the affluent rivers of white and clear water that make up the Amazon basin. Unlike headwater stream piracy events in mountainous terrain — which can produce abrupt barriers such as waterfalls and rapids — the Branco–Negro capture involved tectonic and climatic reorganization of a lowland mega-river system in the Amazonian floodplain [132], where the confluence of the Branco (a clearwater/whitewater river draining the Guiana Shield) with the Negro occurs without elevation discontinuities that would impede large migratory fish. During the transition period, the Rupununi Portal — a vast seasonal wetland (∼3,480 km²) in south-central Guyana that connects upper Branco tributaries to the Essequibo drainage during annual flooding [144,145] — provided an active dispersal stepping stone between the Guiana Shield and Amazon drainages, documented to facilitate directional movement of catfish and other large-bodied freshwater taxa between basins [145]. A further facilitating factor concerns water chemistry: Ruokolainen et al. (2018) [146] documented that the middle and lower Negro basin did not attain its current extreme blackwater conditions (pH 2.9–4.2; conductivity ∼8 µS) until approximately 1,000 years ago, due to the sustained influence of the whitewaterJapurá River, which until recently discharged into the Negro [78,146].

During the Pleistocene colonization window, the lower-middle Negro therefore presented substantially more hospitable physicochemical conditions for *B. rousseauxii* — a large migratory pimelodid dependent on nutrient-rich waters — providing a viable corridor from the upper Branco to the whitewater mainstream Amazon. Consistent with this scenario, Sánchez-Bernal et al. (2023) [78] documented that the cardinal tetra (*Paracheirodon axelrodi*), an obligate blackwater specialist, colonized the Negro River from the Western Guiana Shield via the Cucui corridor during the Late Pleistocene to Holocene (0.255–0.001 Ma) [78], demonstrating the biological viability of this dispersal route for Neotropical fish during this same geological window.

### Disagreeing demographic growth patterns across basins

As indicated above, vicariant/geodispersal effects by uplifting in the Guiana Shield plus climate should have physically separated headwater and generated stream capture. The consequences of this on individual members of the ichthyofauna may cause taxa isolated on either side of new habitats were to experience an effective population size reduction, which may accelerate the genetic divergence between them [147]. Conversely, on the other side of the new habitat, the taxa may use the new connections to colonize and expand their ranges and demographic growth [8]. Here, we observed that although the Orinoco basin was more rapidly and broadly colonized than Amazon, their effective population size resulted in much lower and more stable during a prolonged period (∼1-0.2 Ma) than the effective population size of *B. rousseauxii* in the Amazon.

This Orinoco basin pattern could be explained by its more complex landscape evolution and then, possibly, more critical consequences for its ichthyofaunal establishment and growth. For example, it is known that after subdivision of the Sub-Andean foreland by the rise of the Vaupes Arch, the effects of subsequent marine incursions into the lower Orinoco basin resulted in basin-wide extinctions by reducing the amount of freshwater habitat available to act as a refuge [13,148]. Further, the smaller size of the Orinoco basin may also have contributed to this very-slow population expansion. The size difference between the Orinoco basin (0.88 million km^2^) and Amazon (6.2 million km^2^) is considerable, and therefore the possibility of refuge or availability of resources for *B. rousseauxii* could have been up to 7 times lower for the Orinoco than for the Amazon population. Reduction of the total geographic area and resources available to each new isolated population after a vicariant event, such as that experienced by *B. rousseauxii* on the Guiana Shield, tend to increase rates of extinction or population decline due to landscape subdivision, leading to speciation or extinction as consequence of drift and selection [149,150].

In addition, most vicariant events in South America were accompanied by volcanisms [8,151], being the Orinoco basin one of the most affected. In fact, Lundberg et al. (1998) [10] mentions that extinctions were a common pattern in North South America, suggesting that the causes were related to catastrophic events that characterized the evolutionary history of its ichthyofauna, as it is the last part of the Andes to rise tectonically. This area was characterized by great volcanic activity, especially during the final survey of the Eastern Andes, which took place ∼2.5 Ma [152] , and today hydrologically supports most of the Middle and Lower Orinoco.

Contrary to what happened in the Orinoco with a more delayed population expansion for *B. rousseauxii*, there is evidence that at this same period in the Amazon basin, species such as the piranha *Serrasalmus rhombeus* (∼0.8 Ma), *Colossoma macropomum* (∼0.4 Ma) and *Brachyplatystoma platynemum* (∼0.25 Ma) were experiencing significant growth of their effective population sizes [92,153,154].

On the other hand, it is important to highlight that the most accelerated expansion time in *B. rousseauxii* for both Orinoco and Amazon basins, was within the 150,000 – 130,000 YBP, suggesting that the ecological environment could have been favorable, allowing *B. rousseauxii* occupying more new freshwater area, habitats and adjacent resources for expanding its effective population size.

Interestingly, this rapid expansion coincides with a more moderate temperature and sea levels in the last interglacial, about ca. 130,000– 115,000 YBP [155].

Finally, we can conclude that the separation of the previously continuous Pebas basin into the Orinoco and Amazonas basin by the rise of the Vaupes Arch in the Late Miocene does not explain the distribution and divergence of *B. rousseauxii* in the two basins. Likewise, the black-water Casiquiare canal discovered by Alexander von Humboldt and the Japurá/Guaviare white-water portal in the Andean piedmonts did not serve as a connectivity route between the Orinoco and Amazon populations of *B. rousseauxii*. However, the rivers in the Rupununi Portal of the Guiana Shield, including the recent capture of the Branco River by the Negro River appear to have played a key role in shaping the current patterns of distribution and historical connectivity of *B. rousseauxii* populations of the Orinoco and Amazon basins.

## Acknowledgments

The authors also thank the Laboratório de Evolução e Genética Animal (LEGAL) of the Universidade Federal do Amazonas and the Laboratorio de Ecología de Vertebrados Acuáticos of the Universidad de los Andes for their invaluable scientific support and collaboration throughout this study. This study was part of JGM’s doctoral thesis in the Biotechnology program of UFAM.

## Supporting information

**S1 Fig. Heatmap of the Linkage Disequilibrium index**. Performed in the ‘poppr’ R package [49], it was generated to assess for the existence of linked loci in a matrix of 1738 SNPs (0% missing data) for *Brachyplatystoma rousseauxii* from the Orinoco and Amazon basins (n=30), through the pair.ia() function, calculating the index of association (IA) for all pairs of loci in the dataset. A squared correlation between allelic values at two loci - R2 (rbarD) between 0 and 1 to all pairwise loci comparisons are shown in the analysis.

**S2 Fig. Principal Component Analysis.** The figure shows the quantity of genetic variance explained by the two first components of a Principal Component Analysis for SNPs datasets with 0, 5 and 10% of missing data for *Brachyplatystoma rousseauxii* from the Orinoco and Amazon basins (n=30).

**S3 Fig. The Puechmaille estimators for inferring genetic clusters in the sample**. The figure shows the four Puechmaille estimators (MedMedK, MedMeanK, MaxMedK, and MaxMeanK) supporting K = 2 clusters for the dataset of 12 SSRs loci and 1738 unlinked SNPs from 154 and 30 individuals, respectively, from the gilded catfish *Brachyplatystoma rousseauxii* from the Orinoco and Amazon basins.

**S4 Fig. Bayesian time-calibrated Maximum Credibility Tree**. The figure shows the phylogenetic reconstruction and ancestral relationship of *Brachyplatystoma rousseauxii* individuals from Orinoco and Amazon basins using ddRADseqs and estimated under a GTR+G model of molecular evolution. The observed times are median heights into 95% credibility intervals, calibrated from a nuclear DNA substitution rate of 1.0x10^-9^ mutations per site per year (Freeland, 2005) [74]. Black circles indicate nodes with supports ≥0.95 of posterior probability. The geographical origin of individuals is identified by sample locality IDs in the associated map: Orinoco [Ciudad Guayana (1), Caicara del Orinoco (2), Puerto Carreño (3), Puerto López (4), Guaviare (5), Inírida (6)] and Amazon basin [Boa Vista (7), Manaos (8), Tefé (9), La Pedrera (10)]. River network and background raster data were obtained exclusively from Natural Earth (www.naturalearthdata.com), which provides publicly available geographic datasets licensed under the Creative Commons Attribution 4.0 (CC BY 4.0). No copyrighted or proprietary data sources (e.g., Google Maps, Google Earth, or similar platforms) were used in the creation of this figure. The map was constructed using these datasets within the R statistical environment (www.r-project.org) and edited using Inkscape v1.X (https://inkscape.org).

**S5 Fig. Bayesian time-calibrated Maximum Credibility Tree**. The figure shows the phylogenetic reconstruction and ancestral relationship of *Brachyplatystoma rousseauxii* individuals from Orinoco and Amazon basins using mtDNA concatenated sequences (COI+D-loop) and estimated under an HKY+G model of molecular evolution. The observed times are median heights into 95% credibility intervals, calibrated from a conservative estimate of substitution rate of 0.68x10^-8^ mutations per site per year for the mitochondrial region (Martin & Palumbi 1993) [73]. Black circles indicate nodes with supports ≥0.95 of posterior probability. The geographical origin of individuals is identified by sample locality IDs in the associated map: Orinoco [Ciudad Guayana (1), Caicara del Orinoco (2), Puerto Carreño (3), Puerto López (4), Guaviare (5), Inírida (6)] and Amazon basin [Boa Vista (7), Manaos (8), Tefé (9), La Pedrera (10)]. River network and background raster data were obtained exclusively from Natural Earth (www.naturalearthdata.com), which provides publicly available geographic datasets licensed under the Creative Commons Attribution 4.0 (CC BY 4.0). No copyrighted or proprietary data sources (e.g., Google Maps, Google Earth, or similar platforms) were used in the creation of this figure. The map was constructed using these datasets within the R statistical environment (www.r-project.org) and edited using Inkscape v1.X (https://inkscape.org).

**S1 Table.** Sampling information, final tissue deposit and COI molecular confirmation for *Brachyplatystoma rousseauxii* from Orinoco and Amazon basin.

**S2 Table**. Pairwise population differentiation (*F_ST_*) between sampling localities of *Brachyplatystoma rousseauxii* from the Orinoco and Amazon basins, estimated with microsatellite loci and SNPs.

**S3 Table.** SNP genotype matrix for *Brachyplatystoma rousseauxii* from the Orinoco and Amazon basins containing 1738 loci.

**S4 Table.** Microsatellite genotype matrix for *Brachyplatystoma rousseauxii* from the Orinoco and Amazon basins containing 12 loci.

## Notes

### Competing Interest Statement

The authors have declared no competing interest.

## References

1. Albert JS, Crampton WG. The Geography and Ecology of Diversification in Neotropical Freshwaters. Nature Education Knowledge. 2010;1: 13–19.

2. Kodandaramaiah U. Use of dispersal-vicariance analysis in biogeography - a critique. J Biogeogr. 2009;37: 3–11. doi:10.1111/j.1365-2699.2009.02221.x

3. Lieberman BS. Paleobiogeography: The Relevance of Fossils to Biogeography. Annu Rev Ecol Evol Syst. 2003;34: 51–69. doi:10.1146/annurev.ecolsys.34.121101.153549

4. Reis RE, Albert JS, Di Dario F, Mincarone MM, Petry P, Rocha LA. Fish biodiversity and conservation in South America. J Fish Biol. 2016;89: 12–47. doi:10.1111/jfb.13016

5. Willis SC, Nunes M, Montaña CG, Farias IP, Ortí G, Lovejoy NR. The Casiquiare river acts as a corridor between the Amazonas and Orinoco river basins: biogeographic analysis of the genus Cichla. Mol Ecol. 2010;19: 1014–30. doi:10.1111/j.1365-294X.2010.04540.x

6. Stallard R. River Chemistry, Geology, Geomorphology, and Soils in the Amazon and Orinoco Basins. In: Drever JI, editor. The Chemistry of Weathering. New Jersey: Reidel Publishing Company; 1985. pp. 293– 316.

7. Zeisler R, Ardizzone G. Las aguas continentales de America Latina. 1st ed. FAO Copescal, Tech. Pap. Rome: COPESCAL-FAO; 1979. Available: http://scholar.google.com.secure.sci-hub.org/scholar?q=Las+aguas+continentales+de+Am%C3%A9rica+Latina%2C+de+R.+Ziesler+y+G.D.+Ardizzone%2C+1979&btnG=&hl=es&as_sdt=0%2C5#0

8. Albert JS, Reis RE. Historical biogeography of Neotropical freshwater fishes. 1st ed. Albert JS, Reis RE, editors. Berkely: University of California Press; 2011.

9. Hoorn C, Guerrero J, Sarmiento GA, Lorente MA. Andean tectonics as a cause for changing drainage patterns in Miocene northern South America. Geology. 1995;23: 237–240.

10. Lundberg JG, Marshall LG, Guerrero J, Horton B, Malabarba LMCS, Wesselingh. F. The stage for Neotropical fish diversification: A history of tropical South American rivers. 1st ed. In: Malabarba LR, Reis RE, Vari RP, Lucena ZMS, Lucena CAS, editors. Phylogeny and Classification of Neotropical Fishes. 1st ed. Porto Alegre: Porto Alegre Edipucrs; 1998. pp. 13–48.

11. Winemiller KO, López-Fernández H, Taphorn DC, Nico LG, Duque AB. Fish assemblages of the Casiquiare River, a corridor and zoogeographical filter for dispersal between the Orinoco and Amazon basins. J Biogeogr. 2008;35: 1551–1563. doi:10.1111/j.1365-2699.2008.01917.x

12. Barthem R, Goulding M. The Catfish Connection: Ecology, Migration, and Conservation of Amazon Predators. 1st ed. New York, NY: Columbia University Press; 1997. Available: https://books.google.com/books?hl=es&lr=&id=KbXiUPGREXgC&pgis=1

13. Machado-Allison A. Notas sobre el Origen del Orinoco, su relación con cuencas vecinas, las evidencias biológico-paleotológicas y su conservación: una revisión. Boletín de la Academia de Ciencias Físicas, Matemáticas y Naturales. 2008;68: 25–64.

14. Winemiller KO, Willis SC. The Vaupes Arch and Casiquiare Canal: Barriers and Passages. 1st ed. In: Albert JS, Reis RE, editors. Historical Biogeography of Neotropical Freshwater fishes. 1st ed. Los Angeles: University of California Press; 2011. pp. 225–242.

15. Renza-Millán M, Lasso CA, Morales-Betancourt MA, Villa F, Caballero S. Mitochondrial DNA diversity and population structure of the ocellate freshwater stingray *Potamotrygon motoro* (Müller & Henle, 1841) (Myliobatiformes: Potamotrygonidae) in the Colombian Amazon and Orinoco Basins. Mitochondrial DNA A DNA Mapp Seq Anal. 2019;30: 466–473. doi:10.1080/24701394.2018.1546300

16. Dungel J. Deep in the Jungle. 1st ed. Praga: Euromedia Praga; 2009. Available: http://www.jandungel.com/en/books/po_krk_v_pralese

17. Rice AH. The Rio Negro, the Casiquiare Canal, and the Upper Orinoco, September 1919-April 1920. Geogr J. 1921;58: 321–343. doi:10.2307/1780880

18. Lovejoy NR, Araujo LG. Molecular systematics, biogeography and population structure of Neotropical freshwater needlefishes of the genus Potamorrhaphis. Mol Ecol. 2000;9: 259–268.

19. Escobar MD, Andrade-López J, Farias IP, Hrbek T. Delimiting Evolutionarily Significant Units of the Fish, *Piaractus brachypomus* (Characiformes: Serrasalmidae), from the Orinoco and Amazon River Basins with Insight on Routes of Historical Connectivity. J Hered. 2015;106 Suppl: 428–38. doi:10.1093/jhered/esv047

20. Lovejoy NR, Willis SC, Albert JS. Molecular signatures of Neogene biogeographical events in the Amazon fish fauna. 1st ed. In: Hoorn C, Wesselingh. F, editors. Amazonia: Landscape and Species Evolution: A look into the past. 1st ed. Oxford, UK: Blackwell Publishing; 2011. pp. 405–417. Available: http://onlinelibrary.wiley.com.sci-hub.org/doi/10.1002/9781444306408.ch25/summary

21. Weitzman S, Weitzman M. Biogeography and evolutionary diversification in Neotropical freshwater fishes, with comments on the refuge theory. 1st ed. In: Prance GT, editor. Biological diversification in the tropics. 1st ed. New York, USA: Columbia University Press; 1982. pp. 403–422. Available: http://scholar.google.com.secure.sci-hub.org/scholar?q=Biogeography+and+evolutionary+diversification+in+neotropical+freshwater+fishes%2C+with+comments+on+the+refuge+theory&btnG=&hl=es&as_sdt=0%2C5#0

22. McConnell R, Lowe-McConnell R. Ecological Studides in Tropical Fish communities. Cambridge, UK: Cambridge University Press; 1987. Available: http://scholar.google.com.secure.sci-hub.org/scholar?q=Ecological+Studies+in+Tropical+Fish+Communities&btnG=&hl=es&as_sdt=0%2C5#0

23. Hubert N, Renno J-F. Historical biogeography of South American freshwater fishes. J Biogeogr. 2006;33: 1414–1436. doi:10.1111/j.1365-2699.2006.01518.x

24. Souza LS De, Armbruster J, Werneke D. The influence of the Rupununi portal on distribution of freshwater fish in the Rupununi district, Guyana. Cybium: International journal of ichthyology. 2012;36: 31–43.

25. Lowe-McConnell RH. Speciation in tropical freshwater fishes. Biological Journal of the Linnean Society. 1969;1: 51–75. Available: http://onlinelibrary.wiley.com.sci-hub.org/doi/10.1111/j.1095-8312.1969.tb01812.x/abstract

26. Sabaj M, Taphorn D, Castillo O. Two new species of thicklip thornycats, genus Rhinodoras (Teleostei: Siluriformes: Doradidae). Copeia. 2008; 209–226. Available: http://www.asihcopeiaonline.org.sci-hub.org/perlserv/?request=get-abstract&doi=10.1643%2FCI-05-143

27. Arbour JH, López-Fernández H. Guianacara dacrya, a new species from the rio Branco and Essequibo River drainages of the Guiana Shield (Perciformes: Cichlidae). Neotropical Ichthyology. 2011;9: 87–96. doi:10.1590/S1679-62252011000100006

28. Barthem RB, Goulding M, Leite RG, Cañas C, Forsberg B, Venticinque E, et al. Goliath catfish spawning in the far western Amazon confirmed by the distribution of mature adults, drifting larvae and migrating juveniles. Sci Rep. 2017;7: 1–13. doi:10.1038/srep41784

29. Petrere M, Barthem RB, Córdoba EA, Gómez BC. Review of the large catfish fisheries in the upper Amazon and the stock depletion of piraíba (*Brachyplatystoma filamentosum* Lichtenstein). Reviews in Fish Biology and Fisheries 2005 14:4. 2005;14: 403–414. doi:10.1007/S11160-004-8362-7

30. Barletta M. Estudo da comunidade de peixes bentônicos em tres áreas do canal principal, próximas à confluência dos rios Negro e Solimões, Amazonas (Amazônia Central, Brasil). MSc, Instituto Nacional de Pesquisas da Amazonia, Universidade Federal do Amazonas. 1995. Available: http://www.worldcat.org/title/estudo-da-comunidade-de-peixes-bentonicos-em-tres-areas-do-canal-principal-proximas-a-confluencia-dos-rios-negro-e-solimoes-amazonas-amazonia-central-brasil/oclc/57022090

31. Barthem RB, Fabré NN. Biologia e diversidade dos recursos pesqueiros da Amazônia. In: Ruffinno ML, editor. A pesca e os recursos pesqueiros na Amazonia Projeto manejo dos recursos naturais de Várzea. IBAMA MMA/PROVÁRZEA; 2004. pp. 11–55.

32. Kumar S, Stecher G, Li M, Knyaz C, Tamura K. MEGA X: Molecular Evolutionary Genetics Analysis across computing platforms. Mol Biol Evol. 2018;35(6):1547–1549.

33. Duponchelle F, Pouilly M, Pécheyran C, Hauser M, Renno JF, Panfili J, et al. Sub-regional differentiation of Amazonian catfish (*Brachyplatystoma rousseauxii*, Castelnau, 1855) populations, inferred from otolith microchemical signatures. J Biogeogr. 2016;43: 1020–1033. doi:10.1111/jbi.12690

34. Hauser M, Doria CRC, Melo LFB, Santos AR, Ayala D, Lopes AD, et al. Age and growth of the Amazonian migratory catfish *Brachyplatystoma rousseauxii* in the Madeira River basin before the construction of hydroelectric dams. Neotrop Ichthyol. 2018;16(2): e170122. doi:10.1590/1982-0224-20170122

35. Lundberg JG, Littmann MW. Pimelodidae (Long-whiskered catfishes). In: Reis RE, Kullander SO, Ferraris CJ Jr, editors. Checklist of the Freshwater Fishes of South and Central America. Porto Alegre: EDIPUCRS; 2003. p. 432-446.

36. Ward RD, Zemlak TS, Innes BH, Last PR, Hebert PDN. DNA barcoding Australia’s fish species. Philos Trans R Soc Lond B Biol Sci. 2005;360(1462):1847–1857.

37. Hebert PDN, Cywinska A, Ball SL, deWaard JR. Biological identifications through DNA barcodes. Proc R Soc Lond B. 2003;270(1512):313–321.

38. Ribeiro-Filho DA, Sampaio I, Barros B, Carvalho-Costa LF, Figueiredo KP, Vallinoto M, et al. DNA barcoding reveals mislabeling in Brazil. Food Control. 2020;118:107414.

39. Sambrook J, Fritsch E, Maniatis T. Molecular Cloning: A Laboratory Manual. 2nd ed. New York, NY: Cold Springs Harbor Laboratory Press, Cold Springs Harbor; 1989.

40. Sambrook J, Rusell D. Commonly Used Techniques in Molecular Cloning. 3rd ed. In: Sambrook J, Rusell D, editors. Molecular Cloning. 3rd ed. NY, USA: Cold Spring Harbor Laboratory Press; 2001.

41. Batista JS, Farias IP, Formiga-Aquino K, Sousa a. CB, Alves-Gomes J a. DNA microsatellite markers for “dourada” (*Brachyplatystoma rousseauxii*, Siluriformes: Pimelodidae), a migratory catfish of utmost importance for fisheries in the Amazon: development, characterization and inter-specific amplification. Conserv Genet Resour. 2009;2: 5–10. doi:10.1007/s12686-009-9117-5

42. Martínez JG, Caballero-Gaitán SJ, Sánchez-Bernal D, de Assunção EN, Astolfi-Filho S, Hrbek T, et al. De novo SNP markers development for the Neotropical gilded catfish *Brachyplatystoma rousseauxii* using next-generation sequencing-based genotyping. Conserv Genet Resour. 2016;8. doi:10.1007/s12686-016-0584-1

43. Ivanova N V., Zemlak TS, Hanner RH, Hebert PDN. Universal primer cocktails for fish DNA barcoding. Mol Ecol Notes. 2007;7: 544–548. doi:10.1111/j.1471-8286.2007.01748.x

44. Sivasundar A, Bermingham E, Orti G. Population structure and biogeography of migratory freshwater fishes (*Prochilodus*: Characiformes) in major South American rivers. Mol Ecol. 2001;10: 407–417. doi:10.1046/j.1365-294x.2001.01194.x

45. DeWoody JA, Schupp J, Kenefic L, Busch J, Murfitt L, Keim P. Universal method for producing ROX-labeled size standards suitable for automated genotyping. Biotechniques. 2004;37: 348–352.

46. Peterson BK, Weber JN, Kay EH, Fisher HS, Hoekstra HE. Double digest RADseq: an inexpensive method for de novo SNP discovery and genotyping in model and non-model species. PLoS One. 2012;7: e37135. doi:10.1371/journal.pone.0037135

47. Martínez JG, Rangel-Medrano JD, Yepes-Acevedo AJ, Restrepo-Escobar N, Márquez EJ. Species limits and introgression in *Pimelodus* from the Magdalena-Cauca River basin. Mol Phylogenet Evol. 2022;173. doi:10.1016/j.ympev.2022.107517

48. Kearse M, Moir R, Wilson A, Stones-Havas S, Cheung M, Sturrock S, et al. Geneious Basic: an integrated and extendable desktop software platform for the organization and analysis of sequence data. Bioinformatics. 2012;28: 1647–9. doi:10.1093/bioinformatics/bts199

49. Thompson JD, Higgins DG, Gibson TJ. CLUSTAL W: Improving the sensitivity of progressive multiple sequence alignment through sequence weighting, position-specific gap penalties and weight matrix choice. Nucleic Acids Res. 1994;22: 4673–4680. doi:10.1093/nar/22.22.4673

50. Altschul SF, Gish W, Miller W, Myers EW, Lipman DJ. Basic local alignment search tool. J Mol Biol. 1990;215: 403–410. doi:10.1016/S0022-2836(05)80360-2

51. Martin M. Cutadapt Removes Adapter Sequences From High-Throughput Sequencing Reads. Embnet journal. 2011;17: 10–12. 10.14806/ej.17.1.200

52. Gauthier J, Mouden C, Suchan T, Alvarez N, Arrigo N, Riou C, et al. DiscoSnp-RAD: de novo detection of small variants for RAD-Seq population genomics. PeerJ. 2020;8. doi:10.7717/PEERJ.9291

53. Escobar MD, Barroco L, Martínez JG, Bertuol F, Pouilly M, Freitas CE, et al. How do hydroelectric dams affect non-migratory fish?: genomic evidence for *Cichla temensis* (Perciformes : Cichlidae) in the Uatumã River , Amazonas , Brazil. Biological Journal of the Linnean Society. 2024;143: 1–18.

54. Kamvar ZN, Tabima JF, Grunwald NJ. Poppr: An R package for genetic analysis of populations with clonal, partially clonal, and/or sexual reproduction. PeerJ. 2014;2014: 1–14. doi:10.7717/PEERJ.281/TABLE-6

55. Suzuki M, Ohno K, Sawayama E, Morinaga SI, Kishida T, Matsumoto T, et al. Genomics reveals a genetically isolated population of the Pacific white-sided dolphin (*Lagenorhynchus obliquidens*) distributed in the Sea of Japan. Mol Ecol. 2023;32: 881–891. doi:10.1111/MEC.16797

56. Eaton DAR. PyRAD: Assembly of de novo RADseq loci for phylogenetic analyses. Bioinformatics. 2014;30: 1844–1849. doi:10.1101/001081

57. Van Oosterhout C, Hutchinson WF, Willis DPM, Shipley P. Micro-checker: software for identifying and correcting genotyping errors in microsatellite data. Mol Ecol Notes. 2004;4: 535–538. doi:10.1111/j.1471-8286.2004.00684.x

58. Excoffier L, Lischer HEL. Arlequin suite ver 3.5: a new series of programs to perform population genetics analyses under Linux and Windows. Mol Ecol Resour. 2010;10: 564–7. doi:10.1111/j.1755-0998.2010.02847.x

59. Pritchard JK, Stephens M, Donnelly P. Inference of population structure using multilocus genotype data. Genetics. 2000;155: 945–959.

60. Corander J, Marttinen P, Sirén J, Tang J. Enhanced Bayesian modelling in BAPS software for learning genetic structures of populations. BMC Bioinformatics. 2008;9: 539. doi:10.1186/1471-2105-9-539

61. Earl DA, VonHoldt BM. STRUCTURE HARVESTER: a website and program for visualizing STRUCTURE output and implementing the Evanno method. Conserv Genet Resour. 2011;4: 359–361. doi:10.1007/s12686-011-9548-7

62. Jakobsson M, Rosenberg N a. CLUMPP: a cluster matching and permutation program for dealing with label switching and multimodality in analysis of population structure. Bioinformatics. 2007;23: 1801–6. doi:10.1093/bioinformatics/btm233

63. Rosenberg NA. Distruct: a program for the graphical display of population structure. Mol Ecol Notes. 2004;4: 137–138. doi:10.1046/j.1471-8286.2003.00566.x

64. Evanno G, Regnaut S, Goudet J. Detecting the number of clusters of individuals using the software STRUCTURE: a simulation study. Mol Ecol. 2005;14: 2611–20. doi:10.1111/j.1365-294X.2005.02553.x

65. Puechmaille SJ. The program structure does not reliably recover the correct population structure when sampling is uneven: Subsampling and new estimators alleviate the problem. Mol Ecol Resour. 2016;16: 608–627. doi:10.1111/1755-0998.12512

66. Hey J, Nielsen R. Integration within the Felsenstein equation for improved Markov chain Monte Carlo methods in population genetics. Proceedings of the National Academic of Sciences. 2007;104: 2785– 2790.

67. Putman AI, Carbone I. Challenges in analysis and interpretation of microsatellite data for population genetic studies. Ecol Evol. 2014;4: 4399–428. doi:10.1002/ece3.1305

68. van Oppen MJ, Rico C, Turner GF, Hewitt GM. Extensive homoplasy, nonstepwise mutations, and shared ancestral polymorphism at a complex microsatellite locus in Lake Malawi cichlids. Mol Biol Evol. 2000;17: 489–498. doi:10.1093/oxfordjournals.molbev.a026329

69. Garcia-Vasquez A, Alonso JC, Carvajal F, Moreau J, Nunez J, Renno JF, et al. Life-history characteristics of *Brachyplatystoma rousseauxii* in the Iquitos region, Peru. J Fish Biol. 2009;75(10):2527–2551.

70. Prestes L, Doria CRC, Ruffino ML. Fisheries production of large Amazonian catfish (Siluriformes: Pimelodidae) in the lower Madeira River, before and after the installation of hydroelectric plants. Braz J Biol. 2018;78: 13–22. doi:10.1590/1519-6984.01016

71. Agudelo E, Sánchez CL, Bonilla-Castillo CA, Puerto AM, Valderrama M, Alonso JC, et al. Pesquerías continentales de Colombia: cuencas del Magdalena-Cauca, Sinú, Canalete, Atrato, Orinoco, Amazonas y vertiente del Pacífico. Leticia: SINCHI; 2013.

72. Ricker WE. Computation and interpretation of biological statistics of fish populations. Bull Fish Res Board Can. 1975;191: 1–382.

73. Martin AP, Palumbi SR. Body size, metabolic rate, generation time, and the molecular clock. Proceedings of the National Academy of Sciences. 1993;90: 4087–4091. doi:10.1073/pnas.90.9.4087

74. Freeland JR. Molecular Ecology. 1st ed. eLS. Chichester, England: John Wiley & Sons, Ltd.; 2005. doi:10.1002/9780470015902.a0003268.pub2

75. Hrbek T, Larson A. The evolution of diapause in the killifish family Rivulidae: a molecular phylogenetic and biogeographic perspective. Evolution. 1999;53(4):1200–1216.

76. Jing Y, Long R, Meng J, Yang Y, Li X, Du B, Naeem A, Luo Y. Influence of life-history traits on mitochondrial DNA substitution rates exceeds that of metabolic rates in teleost fishes. Curr Zool. 2025;71(3): 284–294. doi:10.1093/cz/zoae045

77. Ibarra HEA, Segura-Caro JA, Márquez EJ, Martinez JG. Genomic insights into population structure and adaptive variation of *Pimelodus yuma* and *Pimelodus grosskopfii* in the Magdalena-Cauca Basin. PLoS One. 2026;21(6):e0351301. doi:10.1371/journal.pone.0351301

78. Sánchez-Bernal D, Martinez JG, Farias IP, Hrbek T, Caballero S. Phylogeography and population genetic structure of the cardinal tetra (Paracheirodon axelrodi) in the Orinoco basin and Negro River (Amazon basin): evaluating connectivity and historical patterns of diversification. PeerJ. 2023;11: e15117. doi:10.7717/PEERJ.15117/SUPP-12

79. Bergeron LA, Besenbacher S, Zheng J, Liu G, Li P, Bertelsen MF, et al. Evolution of the germline mutation rate across vertebrates. Nature. 2023;615: 285–291. doi:10.1038/s41586-022-05672-7

80. Zhang X, Kivikoski M, Merilä J. De novo mutation rates in sticklebacks. Mol Biol Evol. 2023;40(9): msad192. doi:10.1093/molbev/msad192

81. Kivikoski M, Välimäki K, Merilä J. Rate of de novo mutations in the three-spined stickleback. Heredity. 2025. doi:10.1038/s41437-025-00767-9

82. Drummond AJ, Suchard M a, Xie D, Rambaut A. Bayesian phylogenetics with BEAUti and the BEAST 1.7. Mol Biol Evol. 2012;29: 1969–73. doi:10.1093/molbev/mss075

83. Miller MA, Pfeiffer W, Schwartz T. Creating the CIPRES Science Gateway for inference of large phylogenetic trees. 2010 Gateway Computing Environments Workshop, GCE 2010. 2010. pp. 1–7. doi:10.1109/GCE.2010.5676129

84. Lemey P, Rambaut A, Welch JJ, Suchard MA. Phylogeography takes a relaxed random walk in continuous space and time. Mol Biol Evol. 2010;27: 1877–1885. doi:10.1093/molbev/msq067

85. Posada D. jModelTest: phylogenetic model averaging. Mol Biol Evol. 2008;25: 1253–6. doi:10.1093/molbev/msn083

86. Kates HR, Johnson MG, Gardner EM, Zerega NJC, Wickett NJ. Allele phasing has minimal impact on phylogenetic reconstruction from targeted nuclear gene sequences in a case study of Artocarpus. Am J Bot. 2018;105: 404–416. doi:10.1002/ajb2.1068

87. Baele G, Lemey P, Bedford T, Rambaut A, Suchard MA, Alekseyenko AV. Improving the accuracy of demographic and molecular clock model comparison while accommodating phylogenetic uncertainty. Mol Biol Evol. 2012;29: 2157–2167. doi:10.1093/molbev/mss084

88. Kass R, Raftery A. Bayes’ Factors. J Am Stat Assoc. 1995;90: 773–795. doi:10.1002/9780470015902.a0005851

89. Lemey P, Rambaut A, Drummond AJ, Suchard MA. Bayesian phylogeography finds its roots. PLoS Comput Biol. 2009;5(9):e1000520.

90. Bielejec F, Baele G, Vrancken B, Suchard MA, Rambaut A, Lemey P. SpreaD3: Interactive Visualization of Spatiotemporal History and Trait Evolutionary Processes. Mol Biol Evol. 2016;33: 2167–2169. doi:10.1093/molbev/msw082

91. Bouckaert R, Vaughan TG, Barido-Sottani J, Duchêne S, Fourment M, Gavryushkina A, et al. BEAST 2.5: An advanced software platform for Bayesian evolutionary analysis. PLoS Comput Biol. 2019;15: e1006650. doi:10.1371/journal.pcbi.1006650

92. Farias IP, Torrico JP, García-Dávila C, Santos MDCF, Hrbek T, Renno J-F. Are rapids a barrier for floodplain fishes of the Amazon basin? A demographic study of the keystone floodplain species Colossomamacropomum (Teleostei: Characiformes). Mol Phylogenet Evol. 2010;56: 1129–35. doi:10.1016/j.ympev.2010.03.028

93. Felsenstein J. Accuracy of coalescent likelihood estimates: do we need more sites, more sequences, or more loci? Mol Biol Evol. 2006;23(3):691–700.

94. Lohse K, Barton NH. Estimating divergence parameters with small samples from a large number of loci. Genetics. 2010;184(4):1081–1097.

95. Marandel F, Lorance P, Berthelé O, Trenkel VM, Rivot E, Lamy JB. Estimating effective population size of large marine fish populations using RAD sequencing. Heredity. 2020;125: 326–339. doi:10.1038/s41437-020-00357-x

96. Barthem RB, Ribeiro DB, Lambert MC, Petrere M. Life strategies of some long-distance migratory catfish in relation to hydroelectric dams in the Amazon Basin. Biol Conserv. 1991;55: 339–345. doi:10.1016/0006-3207(91)90037-A

97. Hoorn C. Marine incursions and the influence of Andean tectonics on the Miocene depositional history of northwestern Amazonia: results of a palynostratigraphic study. Palaeogeogr Palaeoclimatol Palaeoecol. 1993;105: 267–309.

98. Hoorn C, Wesselingh F. Amazonia, landscape and species evolution: a look into the past. 1st ed. Oxford, UK: Wiley-Blackwell, John Wiley & Sons; 2011.

99. Albert JS, Val P, Hoorn C. The changing course of the Amazon River in the Neogene : center stage for Neotropical diversification. Neotropical Ichthyology. 2018;16: 1–23. doi:10.1590/1982-0224-20180033

100. Tagliacollo VA, Roxo FF, Duke-Sylvester SM, Oliveira C, Albert JS. Biogeographical signature of river capture events in Amazonian lowlands. J Biogeogr. 2015;42: 2349–2362. doi:10.1111/jbi.12594

101. Lessios HA. The Great American Schism: Divergence of Marine Organisms After the Rise of the Central American Isthmus. Annu Rev Ecol Evol Syst. 2008;39: 63–91. doi:10.1146/annurev.ecolsys.38.091206.095815

102. Turner TF, McPhee M V., Campbell P, Winemiller KO. Phylogeography and intraspecific genetic variation of prochilodontid fishes endemic to rivers of northern South America. J Fish Biol. 2004;64: 186–201. Available: http://onlinelibrary.wiley.com.sci-hub.org/doi/10.1111/j.1095-8649.2004.00299.x/full

103. Hrbek T, Taphorn DC, Thomerson JE. Molecular phylogeny of *Austrofundulus* Myers (Cyprinodontiformes: Rivulidae), with revision of the genus and the description of four new species. Zootaxa. 2005;825: 1–39.

104. Arbogast BS, Kenagy GJ. Comparative phylogeography as an integrative approach to historical biogeography. J Biogeogr. 2001;28(7):819–825.

105. Goulding M. The Fishes and the Forest: Explorations in Amazonian Natural History. 1st ed. Berkely: California University Press; 1980. Available: https://books.google.com/books?hl=es&lr=&id=krIsP5RbFx0C&pgis=1

106. Araujo-Lima CARM, Ruffino ML. Migratory Fishes of South America: Biology, Fisheries and Conservation Status. The Intern. Carolsfeld J, Harvey B, Ross C, Baer A, editors. Ottawa: IDRC/World Bank; 2003. Available: http://www.idrc.ca/EN/Resources/Publications/openebooks/114-0/index.html

107. Sioli H. Studies in Amazonian waters. Golley FB, Medina E, editors. Atas Simposio Biota Amazonica. 1967;3: 9–50.

108. Schmidt GW. Primary production of phytoplankton in the three types of Amazonian waters. III. Primary productivity of phytoplankton in a tropical flood plain lake of Central Amazonia, Lago do Castanho, Amazonas, Brazil. Amazoniana. 1973;4: 379–404. Available: http://web.evolbio.mpg.de/amazoniana/#Amazoniana 4

109. Lewis WM, Hamilton SK, Rodriguez MA, Saunders JF, Lasi MA. Foodweb analysis of the Orinoco floodplain based on production estimates and isotope data. J North Am Benthol Soc. 2001;20: 241–254.

110. Goulding M, Carvalho ML. Life history and management of the tambaqui (Colossoma macropomum, Characidae): an important Amazonian food fish. Rev Bras Zool. 1982;1: 107–133. doi:10.1590/S0101-81751982000200001

111. Morris C, Val AL, Brauner CJ, Wood CM. The physiology of fish in acidic waters rich in dissolved organic carbon, with reference to the Amazon basin. J Exp Zool Part A. 2021;335(9-10):843–863.

112. Gonzalez RJ, Wood CM, Wilson RW, Patrick ML, Val AL. Diverse strategies for ion regulation in fish from the ion-poor, acidic Rio Negro. Physiol Biochem Zool. 2002;75(1):37–47.

113. Lima F, Ribeiro C. Continental-Scale Tectonic Controls of Biogeography and Ecology. 1st ed. In: Albert JS, Reis R, editors. Historical Biogeography Of Neotropical Freshwater Fishes. 1st ed. Berkely: University of California Press; 2011. pp. 145–164. Available: http://drrportal.gov.np/

114. Olivares AM, Hrbek T, Escobar MD, Caballero S. Population structure of the black arowana (Osteoglossum ferreirai) in Brazil and Colombia: Implications for its management. Conservation Genetics. 2013;14: 695– 703. doi:10.1007/s10592-013-0463-1

115. Escalona T, Engstrom TN, Hernandez OE, Bock BC, Vogt RC, Valenzuela N. Population genetics of the endangered South American freshwater turtle, Podocnemis unifilis, inferred from microsatellite DNA data. Conservation Genetics. 2009;10: 1683–1696. doi:10.1007/s10592-008-9746-3

116. Hoorn C. Fluvial palaeoenvironments in the intracratonic Amazonas Basin (early Miocene-early middle Miocene, Colombia). Palaeogeogr Palaeoclimatol Palaeoecol. 1994;109: 1–54. Available: http://www.sciencedirect.com.sci-hub.org/science/article/pii/0031018294901171

117. Martins Formiga K, Da Silva Batista J, Alves-Gomes JA. The most important fishery resource in the Amazon, the migratory catfish *Brachyplatystoma vaillantii* (Siluriformes: Pimelodidae), is composed by an unique and genetically diverse population in the Solimões-Amazonas River System. Neotropical Ichthyology. 2021;19: 2021. doi:10.1590/1982-0224-2020-0082

118. Batista J, Alves-Gomes J. Phylogeography of *Brachyplatystoma rousseauxii* (Siluriformes - Pimelodidae) in the Amazon Basin offers preliminary evidence for the first case of “homing” for an Amazonian migratory catfish. Genetics and Molecular Research. 2006;5: 723–740. Available: http://www.funpecrp.com.br/gmr/year2006/vol4-5/gmr0231_abstract.htm

119. Cionek VDM, Sacramento PA, Zanatta N, Ota RP. Fishes from first order streams of lower Paranapanema and Ivaí rivers, upper Paraná River basin, Paraná, Brazil. Check List. 2012;8: 1158–1162.

120. Diaz-Sarmiento J, Alvarez-León R. Migratory fishes of the Colombian Amazon. 1st ed. In: Carolsfeld J, Harvey B, Ross C, Baer A, editors. Migratory Fishes of South America: Biology, Fisheries and Conservation Status. 1st ed. Ottawa: The World Bank; 2003. p. 384.

121. Montes-Veira S. Cañones de Colombia. 1st ed. Álvarez EO, Castro GS, Aguirre LM, editors. Cali: Banco de Occidente; 2013. Available: http://www.imeditores.com/banocc/canones/

122. Carvajal-Vallejos FM, Duponchelle F, Desmarais E, Cerqueira F, Querouil S, Nuñez J, et al. Genetic structure in the Amazonian catfish *Brachyplatystoma rousseauxii*: influence of life history strategies. Genetica. 2014;142: 323–36. doi:10.1007/s10709-014-9777-2

123. Gravena W, Farias IP, da Silva MNF, da Silva VMF, Hrbek T. Looking to the past and the future: were the Madeira River rapids a geographical barrier to the boto (Cetacea: Iniidae)? Conservation Genetics. 2014;15: 619–629. doi:10.1007/s10592-014-0565-4

124. Souza LS De, Armbruster JW, Willink PW. Connectivity of Neotropical River Basins in the Central Guiana Shield Based on Fish Distributions. Frontiers in Forests and Global Changes. 2020;3: 1–15. doi:10.3389/ffgc.2020.00008

125. Lasso C, Machado-Allison A, Hernández R. Consideraciones zoogeográficas de los peces de La Gran Sabana (Alto Caroni) Venezuela, y sus relaciones con las cuencas vecinas. Memoria de La Sociedad de Ciencias Naturales La Salle. 1990;1990: 133–134. Available: http://scholar.google.com.secure.sci-hub.org/scholar?q=Consideraciones zoogeogr%C3%A1ficas de los peces de la Gran Sabana (Alto Caron%C3%AD) Venezuela%2C y sus relaciones con las cuencas vecinas#0

126. Rosenberg NA, Nordborg M. Genealogical trees, coalescent theory and the analysis of genetic polymorphisms. Nat Rev Genet. 2002;3(5):380–390.

127. Hudson RR, Coyne J. Mathematical consequences of the genealogical species concept. Evolution. 2002;56(8):1557–1565.

128. Degnan JH, Rosenberg NA. Gene tree discordance, phylogenetic inference and the multispecies coalescent. Trends Ecol Evol. 2009;24(6):332–340.

129. Lawson DJ, van Dorp L, Falush D. A tutorial on how not to over-interpret STRUCTURE and ADMIXTURE bar plots. Nat Commun. 2018;9(1):3258.

130. Cremon ÉH, Rossetti D de F, Sawakuchi A de O, Cohen MCL. The role of tectonics and climate in the late Quaternary evolution of a northern Amazonian River. Geomorphology. 2016;271: 22–39. doi:10.1016/J.GEOMORPH.2016.07.030

131. Carolsfeld J, Harvey B, Ross C, Baer A, editors. Migratory Fishes of South America: Biology, Fisheries and Conservation Status. Victoria, BC: World Fisheries Trust/IDRC/World Bank; 2003.

132. Zink RM, Barrowclough GF. Mitochondrial DNA under siege in avian phylogeography. Mol Ecol. 2008;17(9):2107–2121.

133. Toews DPL, Brelsford A. The biogeography of mitochondrial and nuclear discordance in animals. Mol Ecol. 2012;21(16):3907–3930.

134. Ranganathan Y, Karanth A. Conflicting timelines: exploring patterns of mito-nuclear discordance in divergence estimates among tetrapods. 2024. doi:10.1080/10635150.2024.2393437

135. Eytan RI, Hellberg ME. Nuclear and mitochondrial sequence data reveal and conceal different demographic histories and population genetic processes in Caribbean reef fishes. Evolution. 2010;64: 3380–3397. doi:10.1111/j.1558-5646.2010.01071.x

136. Perea S, Vukic J, Sanda R, Doadrio I. Ancient mitochondrial capture promoting mitonuclear discordance in the genus Squalius. PLoS ONE. 2016;11(12):e0166292.

137. Latrubesse EM, Franzinelli E. The late Quaternary evolution of the Negro River, Amazon, Brazil: Implications for island and floodplain formation in large anabranching tropical systems. Geomorphology. 2005;70: 372–397. doi:10.1016/j.geomorph.2005.02.014

138. Wilkinson MJ, Marshall LG, Lundberg JG, Kreslavsky MH. Megafan environments in northern South America. In: Hoorn C, Wesselingh FP, editors. Amazonia: Landscape and Species Evolution. Oxford: Wiley-Blackwell; 2010. pp. 162-184.

139. Fouquet A, Noonan BP, Rodrigues MT, Pech N, Gilles A, Gemmell NJ. Multiple quaternary refugia in the eastern Guiana shield revealed by comparative phylogeography of 12 frog species. Syst Biol. 2012;61: 461–489. doi:10.1093/sysbio/syr130

140. Campbell KE, Frailey CD, Romero-Pittman L. The Pan-Amazonian Ucayali Peneplain, late Neogene sedimentation in Amazonia, and the birth of the modern Amazon River system. Palaeogeogr Palaeoclimatol Palaeoecol. 2006;239: 166–219. doi:10.1016/J.PALAEO.2006.01.020

141. Rossetti DF, Cohen MCL, Tatumi SH, Sawakuchi AO, Cremon ÉH, Mittani JCR, et al. Mid-Late Pleistocene OSL chronology in western Amazonia and implications for the transcontinental Amazon pathway. Sediment Geol. 2015;330: 1–15. doi:10.1016/J.SEDGEO.2015.10.001

142. Cooke GM, Chao NL, Beheregaray LB. Natural selection in the water: freshwater invasion and adaptation by water colour in the Amazonian pufferfish. J Evol Biol. 2012;25: 1305–20. doi:10.1111/j.1420-9101.2012.02514.x

143. Schaefer C, Vale Júnior J. Mudanças climáticas e evolução da paisagem em Roraima: uma resenha do Cretáceo ao recente. 1st ed. In: Barbosa R, Ferreira E, Castellón E, editors. Homem, Ambiente e Ecologia no Estado de Roraima. 1st ed. Manaus, Brazil: Instituto Nacional de Pesquisas da Amazônia, INPA; 1997. pp. 45–64. Available: https://www.passeidireto.com/arquivo/17800242/diversidade-socioambiental-de-roraima/42

144. Lowe-McConnell RH. The fishes of the Rupununi savanna district of British Guiana. Proc Zool Soc Lond. 1964;143(1):121–161.

145. Cardoso de Souza LS, Zuanon J, Leite RG, Torrente-Vilara G, Doria CRC, Dagosta FCP, Julio-Junior HF. Connectivity of Neotropical river basins in the Guiana Shield based on fish distributions. Front For Glob Change. 2020;3:8.

146. Ruokolainen K, Moulatlet GM, Zuquim G, Hoorn C, Tuomisto H. River network rearrangements in Amazonia shake biogeography and civil security. Preprints. 2018. doi:10.20944/preprints201809.0168.v1

147. Templeton AR. The theory of speciation via the founder principle. Genetics. 1980;94: 1011–38. Available: http://www.pubmedcentral.nih.gov/articlerender.fcgi?artid=1214177&tool=pmcentrez&rendertype=abstract

148. Albert JS, Lovejoy NR, Crampton WGR. Miocene tectonism and the separation of cis- and trans-Andean river basins: Evidence from Neotropical fishes. J South Am Earth Sci. 2006;21: 14–27. doi:10.1016/J.JSAMES.2005.07.010

149. Whitlock MC. Fixation probability and time in subdivided populations. Genetics. 2003;164: 767–79. Available: http://www.pubmedcentral.nih.gov/articlerender.fcgi?artid=1462574&tool=pmcentrez&rendertype=abstract

150. Coyne JA, Orr HA. Speciation. 1st ed. Chicago: Sinauer Associates; 2004. Available: https://books.google.com.co/books/about/Speciation.html?id=2Y9rQgAACAAJ&redir_esc=y

151. Schaefer S. The Andes: Riding the Tectonic Uplift. 1st ed. In: Albert J, Reis R, editors. Historical Biogeography of Neotropical Freshwater Fishes. 1st ed. Los Angeles: University of California Press; 2011. pp. 259–278.

152. Hoorn C, Wesselingh FP, Ter Steege H, Bermudez MA, Mora A, Sevink J, et al. Amazonia through time: Andean uplift, climate change, landscape evolution, and biodiversity. Science. 2010;330: 927–931. doi:10.1126/SCIENCE.1194585

153. Hubert N, Duponchelle F, Nuñez J, Rivera R, Bonhomme F, Renno J-F. Isolation by distance and Pleistocene expansion of the lowland populations of the white piranha *Serrasalmus rhombeus*. Mol Ecol. 2007;16: 2488–503. doi:10.1111/j.1365-294X.2007.03338.x

154. Ochoa LE, Pereira LHG, Costa-Silva GJ, Roxo FF, Batista JS, Formiga K, et al. Genetic structure and historical diversification of catfish *Brachyplatystoma platynemum* (Siluriformes: Pimelodidae) in the Amazon basin with implications for its conservation. Ecol Evol. 2015;5: 2005–2020. doi:10.1002/ece3.1486

155. Kopp RE, Simons FJ, Maloof AC, Oppenheimer M. Global and local sea level during the Last Interglacial: A probabilistic assessment. Nature. 2009;462: 863–867. doi:10.1038/nature08686

